# Nonsense Mediated RNA Decay Is a Unique Vulnerability of Cancer Cells with SF3B1 and U2AF1 Mutations

**DOI:** 10.1101/2021.03.19.436079

**Authors:** Abigael Cheruiyot, Shan Li, Sridhar Nonavinkere Srivatsan, Tanzir Ahmed, Yuhao Chen, Delphine Sangotokun Lemacon, Ying Li, Zheng Yang, Brian A. Wadugu, Wayne Warner, Shondra M. Pruett-Miller, Esther A. Obeng, Daniel C. Link, Dalin He, Fei Xiao, Xiaowei Wang, Julie M. Bailis, Matthew J. Walter, Zhongsheng You

## Abstract

Nonsense-mediated RNA decay (NMD) is well recognized as an RNA surveillance pathway that targets aberrant mRNAs with premature translation termination codons (PTCs) for degradation; however, its molecular mechanisms and roles in health and disease remain incompletely understood. In this study, we developed a novel reporter system that can accurately measure NMD activity in individual cells. By carrying out a genome-wide CRISPR/Cas9 knockout screen using this reporter system, we identified novel NMD-promoting factors, including multiple components of the SF3B complex and other U2 spliceosome factors. Interestingly, we also found that cells with mutations in the U2 spliceosome genes *SF3B1* and *U2AF1—*which are commonly found in myelodysplastic syndrome (MDS) and cancers*—*have overall attenuated NMD activity. Furthermore, we found that compared to wild type cells, *SF3B1* and *U2AF1* mutant cells are more sensitive to NMD inhibition, a phenotype that is accompanied by elevated DNA replication obstruction, DNA damage and chromosomal instability. Remarkably, the sensitivity of spliceosome mutant cells to NMD inhibition could be rescued by overexpression of RNase H1, which removes R-loops in the genome. Together, our findings shed new light on the functional interplay between NMD and RNA splicing and suggest a novel strategy for the treatment of MDS and cancers with spliceosome mutations.

## Introduction

In metazoans, pre-mRNA splicing generates diversity in the transcriptome, but it also presents a major source of aberrant RNAs when dysregulated^1,2^. Incorrect splice site selection, intron retention and exon exclusion threaten the fidelity of gene expression, which can cause many genetic disorders, such as β-thalassemia, frontotemporal dementia and laminopathies, and cancer^3–8^. Abnormal splicing is particularly prevalent in myelodysplastic syndrome (MDS) and other cancers with recurring mutations in splicing factors^9,10^. Approximately 50% of MDS, 20% of acute myeloid leukemia (AML) and 60% of chronic myelomonocytic leukemia (CMML) harbor heterozygous somatic mutations in the spliceosome genes *SF3B1*, *U2AF1*, *SRSF2*, and *ZRSR2*, which are involved in the early stage of spliceosome assembly and cause distinct changes in RNA splicing and gene expression^9–21^. Many solid tumors, including uveal melanoma, breast, lung and pancreatic cancers, also harbor spliceosome gene mutations^22–26^. The inherent vulnerability of the splicing process and its dysregulation in disease conditions necessitate mechanisms to detect and control the fate of mis-spliced transcripts. Nonsense-mediated RNA decay (NMD) plays a key role in RNA surveillance by specifically targeting abnormal mRNAs with premature translation termination codons (PTCs) for degradation^27^. NMD also regulates gene expression by degrading physiological transcripts with certain NMD-inducing features, including upstream open reading frames (uORFs), PTC-containing exons, introns in the 3’ untranslated region (UTR), and exceedingly long 3’ UTRs^27–29^. Consequently, NMD modulates the severity of many genetic diseases and regulates various developmental processes and responses to cellular stress^30–36^. In addition to eliminating alternatively spliced or mis-spliced transcripts, NMD is believed to be mechanistically linked to RNA splicing in mammals, as removal of introns from target pre-mRNAs attenuates their degradation by NMD^37–39^. It is believed that splicing-mediated deposition of exon junction complexes (EJCs) facilitates the recognition of PTCs in mRNA, although an EJC-independent NMD pathway also exists^27,40^. A key step of NMD is the recruitment of core NMD factors UPF1 and SMG1 to the terminating ribosome by eRF1 and eRF3, leading to phosphorylation of UPF1 by SMG1, a member of the PIKK family of protein kinases that also include ATM, ATR, DNA-PKcs and mTOR^41,42^. This phosphorylation leads to recruitment of SMG5, SMG6 and SMG7 to target mRNA via phospho-specific interactions, which in turn either directly cleaves the mRNA (SMG6) or recruits nucleolytic activities for RNA degradation (SMG5 and SMG7)^43,44^. Despite extensive research in this area, our understanding of the functional interplay between NMD and splicing remains limited, and the exact role of NMD in cells with defective splicing remains to be determined.

In this study, we have identified multiple early splicing factors that promote NMD, and a synthetic lethal relationship between splicing dysregulation and NMD disruption. By performing a genome-wide CRISPR/Cas9 knockout screen using a novel NMD reporter system, we have identified a number of new factors that promote NMD, including components of the U2 spliceosome complex. Interestingly, cells expressing mutants of the U2 spliceosome genes *SF3B1* or *U2AF1* that are frequently found in MDS and cancers exhibited attenuated NMD activity. Furthermore, we found that spliceosome mutant cells are hypersensitive to NMD disruption. Remarkably, this sensitivity could be rescued by ectopic expression of RNaseH1, which removes R loops, a cellular structure containing a RNA/DNA hybrid and displaced ssDNA^45–47^. Together our results have uncovered novel factors that promote NMD and a new NMD-targeting strategy for treating MDS and other cancers with defective splicing.

## Results

### A novel reporter system that measures NMD activity in individual cells

In order to explore the mechanisms and functional regulation of the NMD pathway, we sought to identify additional NMD factors and regulators through a genome-wide CRISPR/Cas9 screen. To this end, we developed a new reporter system that can rapidly and accurately measure NMD activity in individual mammalian cells. This new reporter system (Fig. 1A) is built on a bioluminescence-based NMD reporter that we developed previously, which consists of two separate, but highly homologous transcription units that are inserted in tandem into a single vector^35,48^. Each unit in the original reporter contains a CMV promoter, a T cell receptor-β (TCRβ) minigene containing three exons and two introns, a HA tag-encoding sequence inserted in exon 1, and a polyadenylation signal. The first unit, which contains the open reading frame (ORF) of the CBR luciferase gene and its natural stop codon in exon 2 of the TCRβ minigene, expresses a nonsense mRNA that is targeted for degradation by NMD. The second unit, which serves as an internal control for the expression of the first unit, contains the ORF of the CBG99 luciferase gene (without a stop codon) in the same position in exon 2 of the TCRβ minigene. This original reporter can be used to measure NMD activity in a population of cells, based on the ratio of the products of the two fusion reporter genes at the levels of RNA, protein, or the luciferase activity of CBR and CBG^35,48,49^. To develop a reporter that can analyze NMD activity in individual cells, we inserted the ORFs of mCherry and EGFP (without stop codons) immediately upstream of CBR and CBG, respectively, into the original reporter (Fig. 1A). The increase and decrease in the mCherry/EGFP signal ratio represent NMD repression and enhancement, respectively. Both the fluorescent proteins (mCherry and EGFP) and luciferases (CBR and CBG) in the reporter are functional (Figure 1). Thus, this new reporter is expected to allow for accurate NMD analysis in individual live cells through fluorescence detection, while still retaining the ability to measure NMD efficiency in a group of cells via bioluminescence detection.

**Figure 1.**
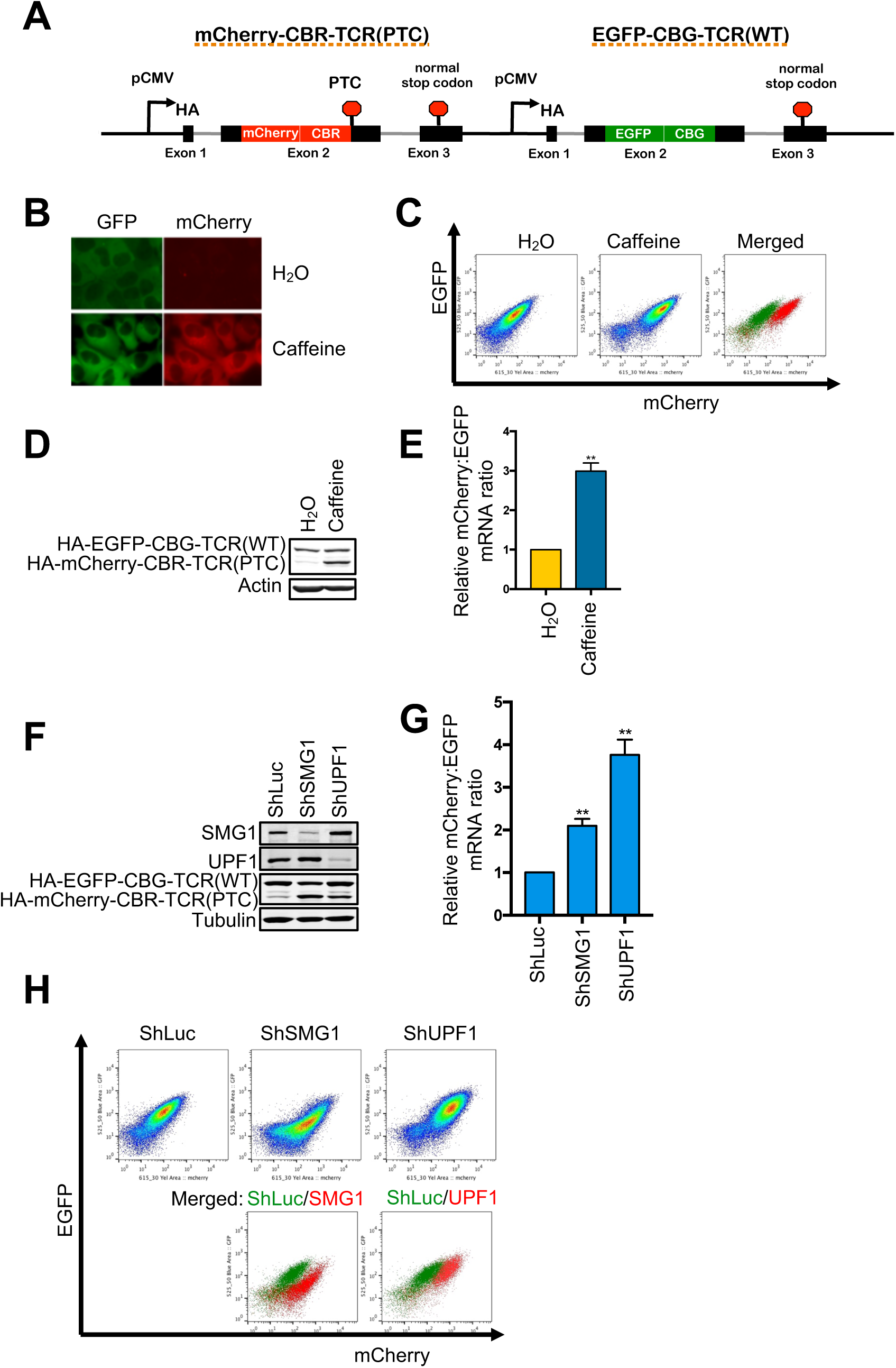
A new reporter system for analyzing NMD in individual human cells. A. Schematic diagram of a multicolored, fluorescence- and bioluminescence-based NMD reporter. B. Fluorescence imaging of the NMD reporter in U2OS reporter cells after treatment with H_2_O or caffeine (10 mM) for 24 hrs. C. FACS analysis of U2OS reporter cells treated with H_2_O or caffeine (10 mM) for 24 hrs. In the merged panel, green dots are H_2_O-treated cells whereas red dots are caffeine-treated cells. D. Western blot of the protein products (HA-tagged) of the NMD reporter after 24-hr treatment of U2OS reporter cells with H_2_O or caffeine (10 mM). E. Ratios of mCherry-containing reporter mRNA to EGFP-containing reporter mRNA in U2OS reporter cells treated with H_2_O or caffeine (10 mM) for 24 hrs. The mCherry/EGFP mRNA ratio of the H_2_O control was normalized to 1. Data represent the mean ± SD of three independent experiments. **p ≤ 0.01 (paired t-test). F. Western blot of the protein products of the NMD reporter in U2OS reporter cells after shRNA-mediated knockdown of SMG1 or UPF1. G. Ratio of mCherry-containing reporter mRNA to EGFP-containing reporter mRNA in U2OS reporter cells after shRNA-mediated knockdown of SMG1 or UPF1. The mCherry/EGFP mRNA ratio of the shLuc control was normalized to 1. Data represent the mean ± SD of three independent experiments. **p ≤ 0.01 (paired t-test). H. FACS analysis of U2OS reporter cells after shRNA-mediated knockdown of SMG1 or UPF1. In the merged figure, green dots represent control shLuc cells, and red dots represent shSMG1 or shUPF1 samples.

To validate the new NMD reporter, we generated a U2OS cell line stably expressing the reporter (hereafter referred to as U2OS reporter cells) through stable transfection and clone validation. As expected, while the reporter cells exhibited robust EGFP signal, little mCherry signal was detected by fluorescence imaging (Fig. 1B). Treatment of the reporter cells with caffeine, which inhibits NMD by decreasing the enzymatic activity of the SMG1 protein kinase, increased the mCherry signal (Fig. 1B). Flow cytometry analysis also showed an increase in the mCherry/EGFP fluorescence ratio after caffeine treatment, leading to a shift of the cell population in a dot plot (Fig. 1C). These results were corroborated by western blot and RT-qPCR analyses of the levels of protein and RNA, respectively, of the two fusion reporter genes (Fig. 1D and E). Additionally, shRNA- or sgRNA-mediated depletion of SMG1, UPF1, or UPF2 also resulted in increased mCherry/EGFP ratio at the levels of protein, RNA and fluorescence activity (Fig. 1F-H, Suppl. Fig. 1A-C), further validating the reporter. Together, these data demonstrate that our new reporter is a specific, robust and convenient system for analyzing NMD activity at both single-cell and population levels.

### A genome-wide CRISPR/Cas9 knockout screen identifies novel NMD-promoting factors in human cells

Using the new NMD reporter system described above, we next performed a genome-wide CRISPR/Cas9 knockout screen to identify new NMD factors and regulators. To do this, we generated a U2OS reporter cell line expressing Cas9 and infected the cells with a lentiviral GeCKOv2 human sgRNA library to knock out individual genes in cells. Fluorescence-activated cell sorting (FACS) was then performed to collect cells with inhibited NMD activity (0.22% of infected cells with increased mCherry/EGFP ratio) (Fig. 2A and B). Genomic DNA was then isolated from the collected cells as well as a fraction of infected but unsorted cells (baseline control), and the integrated sgRNA inserts in the genome were amplified by PCR. After the addition of Illumina sequencing tags via a PCR method, samples were subjected to Next-Gen sequencing and analysis to obtain read counts for each sgRNA in the library. MAGeCK analysis was then performed to rank genes based on the enrichment of their respective sgRNAs in the collected NMD-inhibited cells. Notably, among the 15 top ranked hits, six are known to promote NMD. These include three known NMD factors (*UPF1*, *SMG6*, *RUVBL1*), two components of the EJC (*eIF4A3*, *RBM8A*), and a spliceosome factor that facilitates EJC assembly on mRNA (*CWC22*) (Fig. 2C and D, Suppl. Table 3). Moreover, Gene Set Enrichment Analysis (GSEA) of the overall screen result indicates that NMD, spliceosome and mRNA translation are among the most enriched pathways (Fig. 2E). Together, these data further validate our new reporter system and the quality of the genome-wide CRISPR/Cas9 screen.

**Figure 2.**
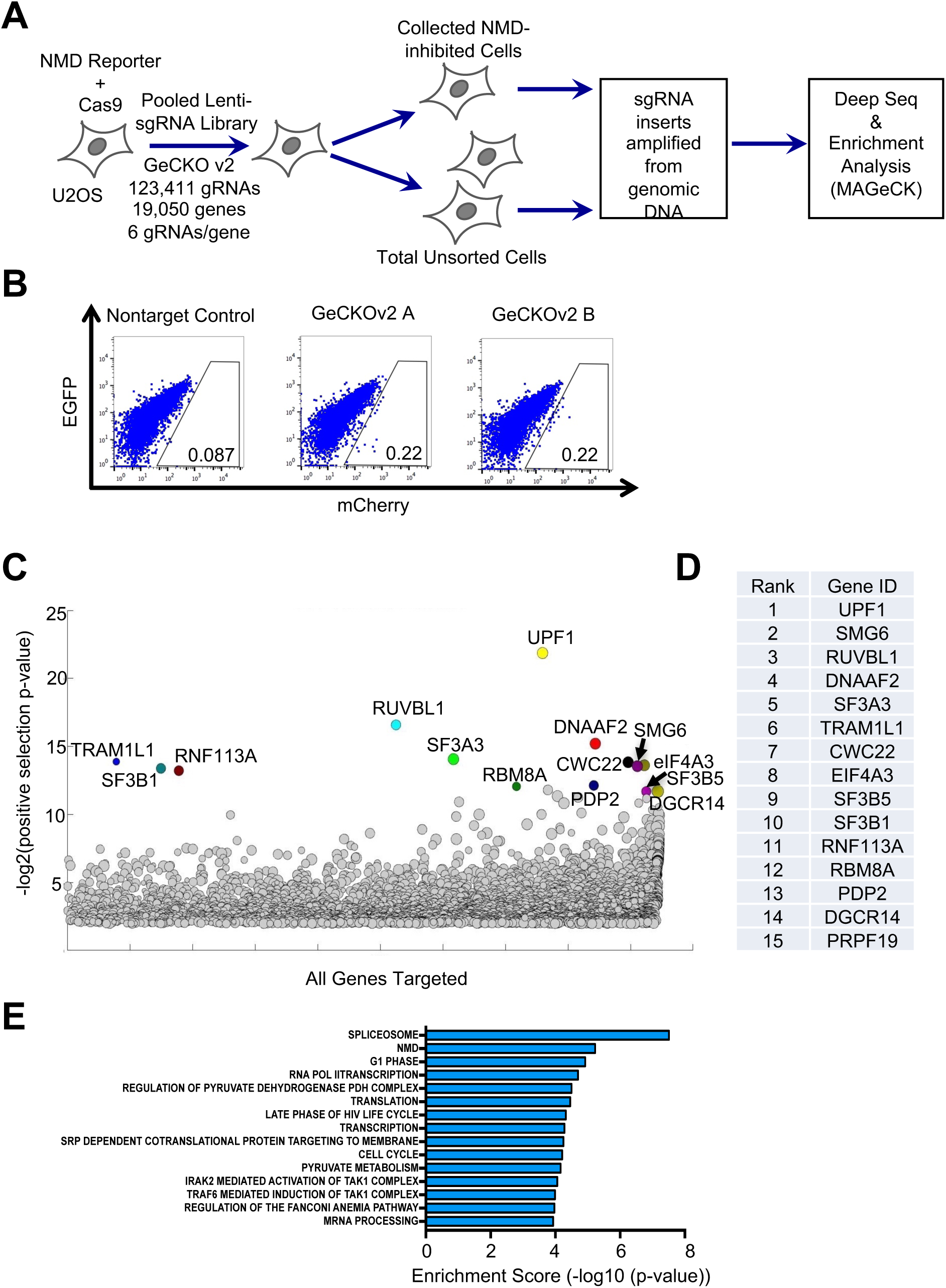
A genome-wide CRISPR/Cas9 knockout screen to identify novel NMD factors and regulators. A. Workflow of the CRISPR/Cas9 knockout screen. B. FACS analysis of Cas9-expressing U2OS reporter cells infected with the two GeCKOv2 sgRNA sub-libraries, or a non-targeting sgRNA control. The gating represents the collected cell population with attenuated NMD activity. C. A bubble plot showing results of gene enrichment analysis obtained from MAGeCK analysis. The bubble size represents the number of gRNAs enriched for the target gene. D. The list of top 15 gene hits as ranked by MAGeCK analysis. E. GSEA analysis of the ranked gene list.

In addition to these known NMD factors, our screen identified several genes that are potentially novel NMD-promoting factors in human cells, providing a rich resource for future elucidation of the mechanisms and regulation of the NMD pathway. In the present study, we validated 9 genes among the top 15 hits that were not known to be involved in NMD. These genes are involved in the early stage of spliceosome assembly during splicing (*SF3B1*, *SF3B5*, *SF3A3*, *PRPF19*, *RNF113A*, *DGCR14*)^50–53^, regulation of pyruvate dehydrogenase activity (*PDP2*)^54^, cilia function (*DNAAF2*)^55^, or unknown processes (*TRAM1L1*). Using two independent sgRNAs for each gene that are distinct from that in the original GeCKOv2 library, we examined the effects of knockdown of these factors on NMD of our reporter. Western blot results show that depletion of each factor increased the expression of the HA-mCherry-CBR-TCR(PTC) fusion protein as well as the HA-mCherry-CBR-TCR(PTC)/HA-EGFP-CBG-TCR(WT) ratio (Suppl. Fig. 2A and B), suggesting that these factors indeed promote NMD. To complement this experiment, we examined in Calu-6 cells the effect of depletion of these factors on the stability of p53 mRNA, which contains an endogenous PTC and is known to be degraded by NMD^41,56^. Depletion of the aforementioned 9 factors individually caused increased stability of p53 mutant mRNA, consistent with the results of our reporter assay (Supplementary Fig. 2C). Furthermore, depletion of these factors also increased the stability of several physiological NMD targets in Calu-6 cells, including *ATF4*, *PIM3*, and *UPP1*, but not the stability of *ORCL*, which is not a NMD target (Suppl. Fig. 2D-G)^56–58^. These observations independently verify our CRISPR screen results, although further characterization is needed to define the exact roles of these genes in NMD.

### NMD activity is attenuated in cells with SF3B1 and U2AF1 mutations

Our genome-wide CRISPR/Cas9 knockout screen identified the U2 spliceosome genes *SF3B1*, *SF3B5*, and *SF3A3* among the top candidate genes that promote NMD. Interestingly, a recent siRNA-based genome-wide screen by Baird et al^59^ using a different NMD reporter system also identified U2 spliceosome genes, including *U2AF1* and SF3 complex genes, among the top hits that promote NMD. The enrichment of U2 spliceosome factors in the results of two completely different screens raises the possibility that these factors play a unique role in NMD in addition to their function in RNA splicing. Interestingly, the U2 genes *U2AF1* and *SF3B1*, as well as U2/U12-related gene *ZRSR2* are also frequently mutated in MDS and cancers, and these mutations cause aberrant splicing^9–17,19^. The identification of multiple U2 spliceosome genes as putative NMD factors/regulators in at least two genome-wide screens motivated us to determine whether common MDS-associated *U2AF1* and *SF3B1* mutations also affect NMD activity.

Because both U2AF1 and SF3B1 are required for RNA splicing, which is mechanistically linked to NMD, we examined the effects of mutations of these spliceosome factors on NMD by using a NMD reporter mRNA that is not spliced. We generated an intronless, tethering reporter system for NMD analysis, based on a reporter system previously developed by Gehring et al^60^. The published system consists of a transcription unit encoding β-globin reporter pre-mRNA with four boxB sites in the 3’ UTR, and a λN-fused NMD factor whose binding to boxB sites leads to degradation of the target mRNA through NMD^60–64^. Building on this, we generated an EGFP-boxB reporter by replacing the β-globin reporter pre-mRNA sequence with the EGFP ORF (Fig. 3A). We then generated a U2OS cell line stably expressing this reporter. As expected, no cryptic splicing was detected in this intronless reporter mRNA in cells (Suppl. Fig. 3A). As a control, we also generated a U2OS cell line stably expressing a reporter mRNA with scrambled boxB sites (boxB’) in the 3’ UTR that are deficient in λN binding. Expression of λN fusion proteins in cells was controlled by a doxycycline-inducible system. To assess the stability of the reporter transcripts, actinomycin D was used to block transcription, and the amount of reporter mRNA remaining before and after actinomycin D treatment was measured by RT-qPCR. Consistent with published results, expression of λN-fused UPF3B (a known NMD factor) (λN-UPF3B) resulted in accelerated degradation of the EGFP-boxB reporter mRNA, while expression of λN or UPF3B alone had no effect (Fig. 3B). Expression of λN, UPF3B or λN-UPF3B did not affect the stability of the EGFP-boxB’ reporter transcript (Fig. 3B). Importantly, the degradation of the EGFP-boxB reporter mRNA after λN-UPF3B induction was completely abrogated by sgRNA/Cas9-mediated depletion of the key NMD factor UPF1 (Fig. 3C). Treating cells with a specific small molecule inhibitor of SMG1 (SMG1i), which does not inhibit related kinases ATR, mTOR and PI3K at the used concentrations (Suppl. Fig. 3B-E), also prevented the degradation of the EGFP-boxB reporter mRNA (Fig. 3D). Together, these results further validate the degradation of the λN-UPF3B-tethered EGFP-boxB reporter mRNA by NMD.

**Figure 3.**
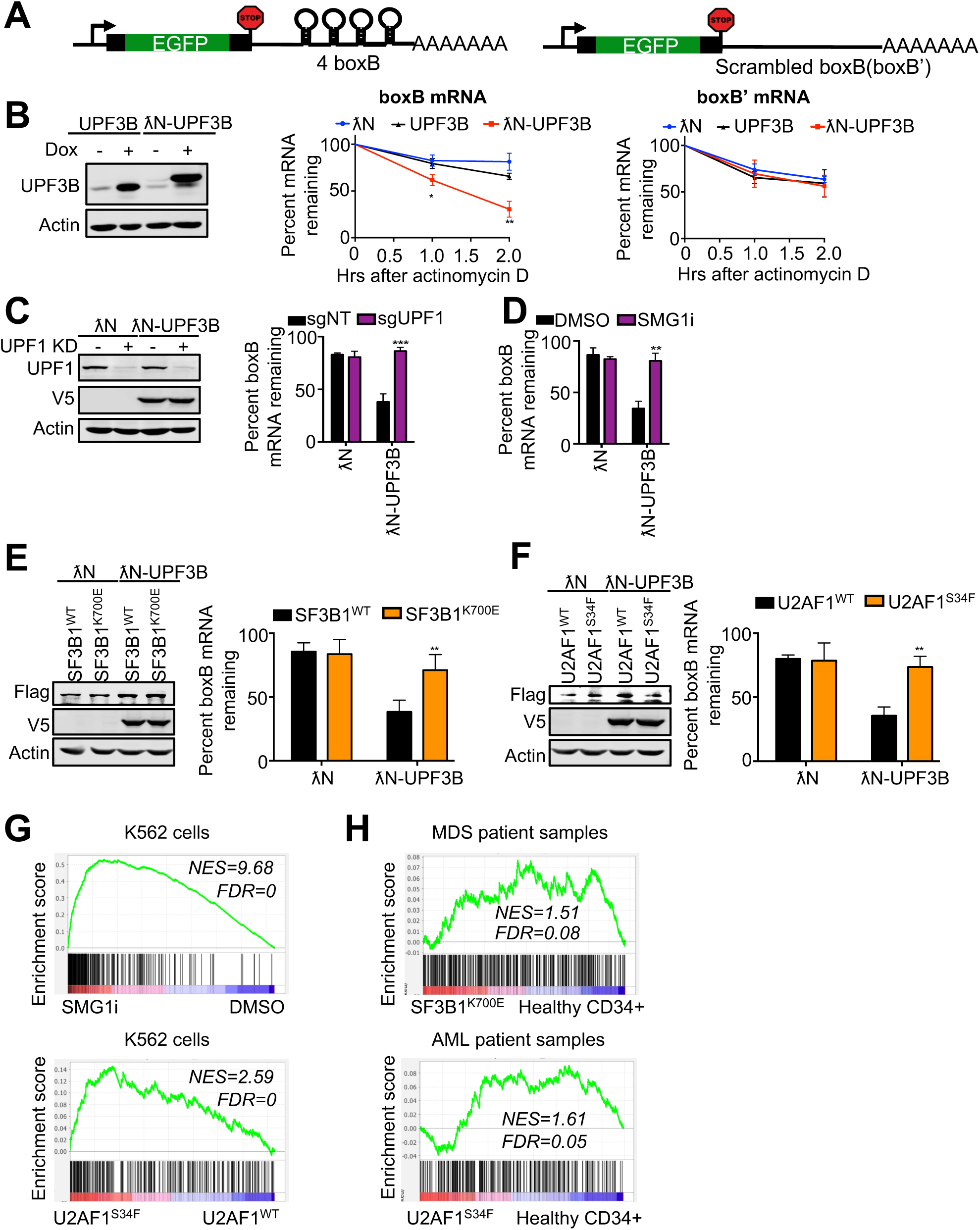
NMD activity is attenuated in cells with SF3B1 and U2AF1 mutations. A. A schematic of a tethering reporter that recapitulates NMD in human cells. The 3’ UTR of the reporter construct contains 4 boxB sites. A reporter with scrambled boxB (boxB’) sequences in the 3’ UTR was used as a control. B. Left, western blot analysis of UPF3B or λN-UPF3B proteins (both V5-tagged) after induction with doxycycline (1 ug/ml). Right, stability of EGFP-boxB or EGFP-boxB’ reporter mRNA in cells expressing λN, UPF3B, or λN-UPF3B. RNA decay analysis was performed by measuring RNA before and after actinomycin D treatment. Percent mRNA remaining was calculated as the mRNA remaining as a percent of RNA before actinomycin D treatment. Data represent the mean ± SD of three independent experiments. **p ≤ 0.01, *p ≤ 0.05 (unpaired t-test). C. Left, western blot analysis of UPF1 knockdown and λN-UPF3B induction in cells expressing the boxB reporter mRNA. Right, effects of UPF1 knockdown on the stability of EGFP-boxB reporter mRNA in cells expressing λN or λN-UPF3B. Data represent the mean ± SD of three independent experiments. ***p ≤ 0.001 (unpaired t-test). D. RNA decay analysis of EGFP-boxB reporter mRNA in λN- or λN-UPF3B expressing cells after treatment with DMSO or SMG1i (1 μM). Data represent the mean ± SD of three independent experiments. **p ≤ 0.01 (unpaired t-test). E-F. Left, western blot analysis of λN-UPF3B in EGFP-boxB reporter cells expressing Flag-tagged SF3B1^WT/K700E^ (E) or U2AF1^WT/S34F^ (F). Right, effects of SF3B1^WT/K700E^ or U2AF1^WT/S34F^ overexpression on the stability of EGFP-boxB reporter mRNA in cells in the presence of λN-UPF3B. Data represent the mean ± SD of three independent experiments. **p ≤ 0.01 (unpaired t-test). G. Gene Set Enrichment Analysis (GSEA) enrichment score plots for NMD target genes that are upregulated by UPF1 KD. RNA was isolated from K562 cells and sequenced to determine the effects of SMG1i treatment (1 μM for 3 days, top) or U2AF1^S34F^ expression (bottom) on gene expression. Individual genes in the gene set are represented by a black vertical bar at the bottom of the plot. H. Gene Set Enrichment Analysis (GSEA) enrichment score plots for NMD target genes that are upregulated by UPF1 KD. RNA was isolated from AML, MDS or control samples and sequenced to determine the effects of SF3B1^K700E^ expression (top, analysis of MDS data in Pellagatti et al.^121^) and U2AF1^S34F^ expression (bottom, AML samples) on gene expression. Individual genes in the gene set are represented by a black vertical bar at the bottom of the plot.

To determine whether *SF3B1* and *U2AF1* mutations affect NMD activity, we examined the effects of *SF3B1^K700E^* or *U2AF1^S34F^*, both of which are commonly found in MDS and cancers, on the decay of λN-UPF3B-tethered EGFP-boxB reporter mRNA by NMD. To do this, we expressed Flag-SF3B1^K700E^, Flag-SF3B1^WT^, Flag-U2AF1^S34F^, or Flag-U2AF1^WT^ in U2OS cells that contain the tethering reporter system for NMD described above. The effects on NMD of the EGFP-boxB reporter mRNA after λN-UPF3B induction were assessed via RT-qPCR. Interestingly, as shown in Fig. 3E-F, cells expressing Flag-SF3B1^K700E^ or Flag-U2AF1^S34F^, but not Flag-SF3B1^WT^ or Flag-U2AF1^WT^, exhibited a reduced level of degradation of the reporter RNA, suggesting that NMD activity is attenuated in the presence of these spliceosome mutants.

To further corroborate these results, we sought to determine whether expression of these spliceosome mutants in cells increases the levels of physiological NMD targets as would be predicted from reduced NMD activity. To this end, we created a custom list of putative endogenous NMD targets based on the genes identified by Longman et al^65^ that were upregulated after UPF1 knockdown. As expected, enrichment of upregulated putative NMD targets was observed in K562 cells treated with the NMD inhibitor SMG1i (Fig. 3G, top panel). Remarkably, GSEA analysis of mRNAs in K562 cells expressing U2AF1^S34F^ showed that these spliceosome mutations also upregulated putative physiological NMD targets (Fig. 3G, bottom panel). Similarly, AML and MDS patient samples harboring U2AF1^S34F^ and SF3B1^K700E^ mutations, respectively, also exhibited enrichment of upregulated putative endogenous NMD targets (Fig. 3H). These results support the idea that *SF3B1* and *U2AF1* mutations attenuate NMD activity. To further test this idea, we created another custom list of putative physiological NMD targets based on the RNA-seq results generated in this study on the effects of SMG1i treatment on gene expression. Consistent with the results for the UPF1 depletion-based gene-set, the expression of U2AF1^S34F^ or SF3B1^K700E^ also caused a strong increase in the expression of the SMG1i-based gene-set of putative NMD targets (Supp. Fig. 3F). Together, results of our reporter assay and endogenous NMD target analysis strongly suggest that MDS-associated mutations in *SF3B1* and *U2AF1* attenuate NMD.

### Cancer cells harboring spliceosome mutations are preferentially sensitive to NMD attenuation

The high levels of nonsense mRNAs observed in cells with spliceosome mutations and the role of NMD in the clearance of these potentially deleterious transcripts raise the possibility that these mutant cells are dependent on NMD for survival^10,17,18^. The attenuated NMD activity in spliceosome mutant cells may also make them more vulnerable to further NMD disruption. In support of this idea, we found that SF3B1^K700E^-expressing U2OS cells exhibited much more reduced viability after shRNA-mediated knockdown of UPF1, compared to cells expressing a comparable level of SF3B1^WT^ (Fig. 4A). Similarly, U2OS cells expressing U2AF1^S34F^ were also more sensitive to UPF1 knockdown than cells expressing U2AF1^WT^ (Fig. 4B). These results suggest that a synthetic lethal relationship exists between spliceosome mutations and NMD disruption and that NMD can be targeted to selectively eliminate spliceosome mutant cells. In further support of this idea, we found that SF3B1^K700E^-expressing U2OS cells were much more sensitive to SMG1i, compared to SF3B1^WT^-expressing cells (Fig. 4C). K562 leukemia cells expressing U2AF1^S34F^ also displayed heightened sensitivity to SMG1i, compared to cells expressing U2AF1^WT^ (Fig. 4D). Furthermore, SMG1i treatment preferentially killed CRISPR/Cas9-engineered K562 cells that contained a SF3B1^K666N^ knock-in mutation, compared to the isogenic wild type control cells (Fig. 4E, Suppl. Fig 4A). Together these data suggest the possibility that NMD is a therapeutic vulnerability for cancer cells with spliceosome mutations. It was previously reported that spliceosome mutant cells are sensitive to splicing modulators such as PB, sudemycin, E7107, and H3B-8800^18,66–68^. Consistent with published results, U2OS or K562 cells expressing SF3B1^K700E^, U2AF1^S34F^, or SF3B1^K666N^ all exhibited elevated sensitivity to PB (Fig. 4C-E).

**Figure 4.**
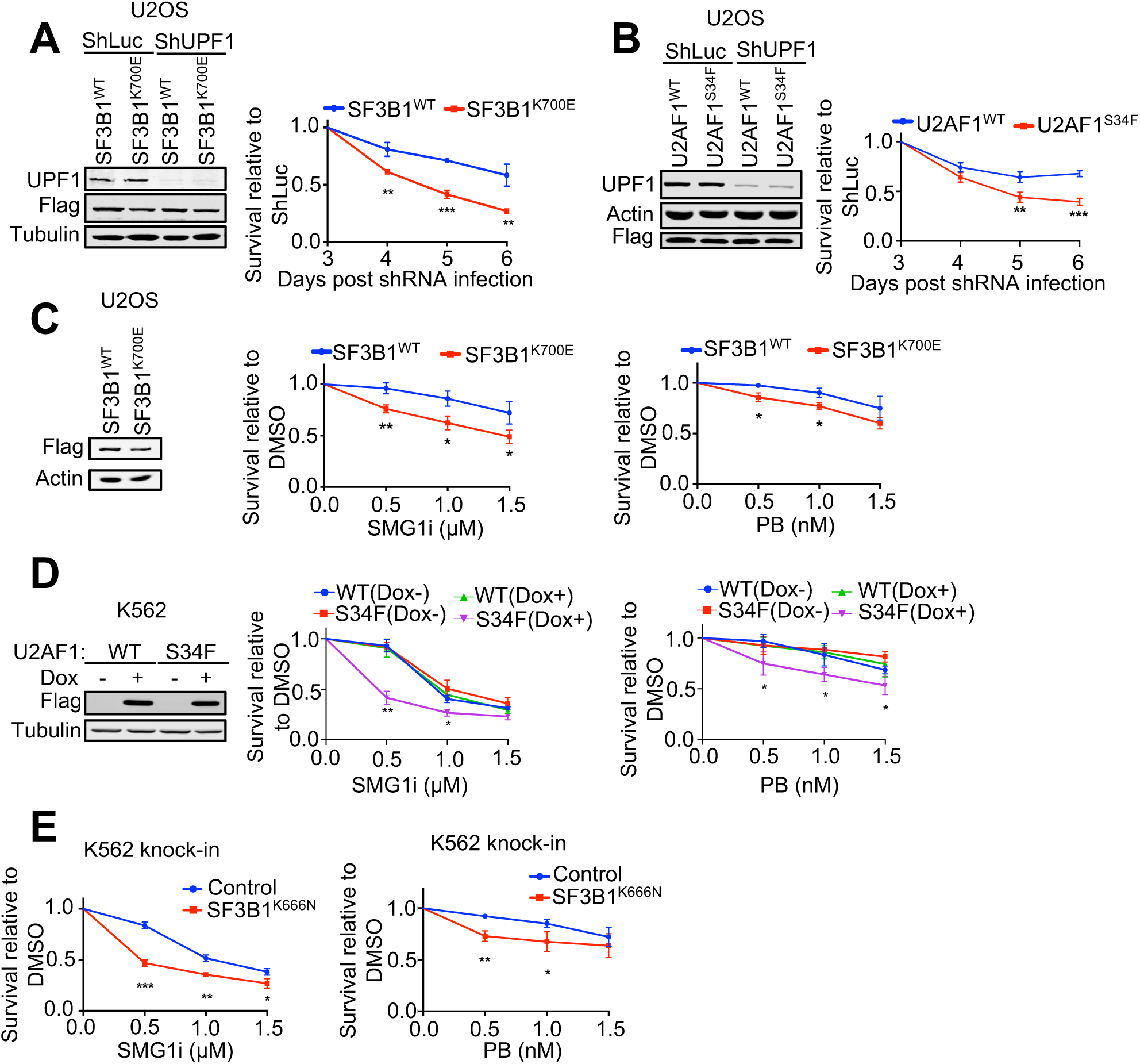
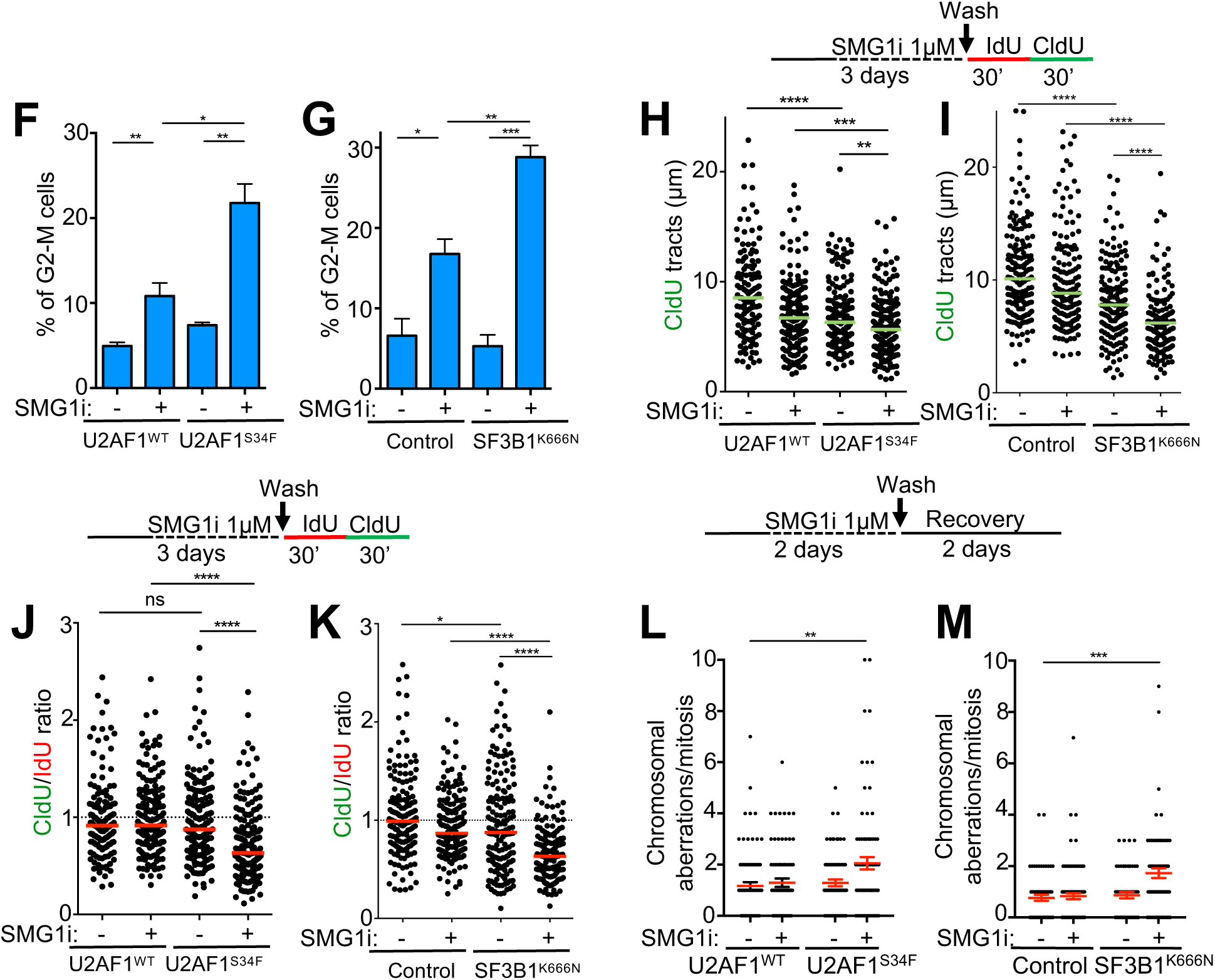
Cells expressing mutant spliceosome factors are sensitive to NMD inhibition. A-B. Left, western blot analysis of UPF1 knockdown in U2OS cells expressing Flag-tagged SF3B1^WT/K700E^ (A) or U2AF1^WT/S34F^ (B). Right, effects of UPF1 knockdown on the viability of U2OS cells expressing SF3B1^WT/K700E^ (A) or U2AF1^WT/S34F^ (B). Data represent the mean ± SD of three independent experiments. ***p ≤ 0.001; **p ≤ 0.01 (unpaired t-test). C. Left, western blot analysis of Flag-tagged SF3B1^WT^ or SF3B1^K700E^ overexpression in U2OS cells. Middle and right, effects of SMG1i or PB treatment (3 days) on the viability of U2OS cells expressing SF3B1^WT^ or SF3B1^K700E^. Data represent the mean ± SD of three independent experiments. **p ≤ 0.01; *p ≤ 0.05 (unpaired t-test). D. Left, western blot analysis of inducible expression of Flag-tagged U2AF1^WT^ or U2AF1^S34F^ in K562 cells. Middle and right, effects of SMG1i or PB treatment (3 days) on the viability of K562 cells expressing U2AF1^WT^ or U2AF1^S34F^. Data represent the mean ± SD of three independent experiments. **p ≤ 0.01; *p ≤ 0.05 (unpaired t-test). E. Effects of SMG1i or PB treatment (3 days) on the viability of K562 cells with or without SF3B1^K666N^ knock-in mutation. Data represent the mean ± SD of three independent experiments. ***p ≤ 0.001; **p ≤ 0.01; *p ≤ 0.05 (unpaired t-test). F-G. Effects of SMG1i treatment (1 μM, 3 days) on the G2-M population of the K562 cells expressing U2AF1^WT/S34F^ (F) or K562 cells with or without SF3B1^K666N^ knock-in mutation (G). Data represent the mean ± S.E.M of three independent experiments. ***p ≤ 0.001; **p ≤ 0.01; *p ≤ 0.05 (unpaired t-test). H-K. Effects of SMG1i treatment (1 μM, 3 days) on DNA replication speed (H, I) or fork progression (J, K) in K562 cells expressing U2AF1^WT/S34F^ (H, J) or K562 cells with or without SF3B1^K666N^ knock-in mutation (I, K). Upper, experimental scheme for DNA fiber assay. Lower, distribution of CldU tract lengths or CIdU/IdU ratio. Green or red bars represent the median ± S.E.M of two independent experiments. A total of 150 tracts analyzed per sample. ****p ≤ 0.0001; ***p ≤ 0.001; **p ≤ 0.01 (Mann Whitney test). L-M. Effects of SMG1i treatment (1 μM, 2 days followed by 2 days of recovery) on chromosomal integrity in K562 cells expressing U2AF1^WT/S34F^ (L) or K562 cells with or without SF3B1^K666N^ knock-in mutation (M). Upper, experimental scheme. Lower, distribution of chromosomal aberrations per mitosis. Red bars represent the mean ± S.E.M of two independent experiments. A total of 150 metaphases analyzed per sample. **p ≤ 0.01 (Mann Whitney test).

To explore the cellular processes responsible for the sensitivity of spliceosome mutant cells to NMD inhibition, we first examined the effects of SMG1i treatment on cell cycle progression and DNA replication. To do this, we performed flow cytometry analysis on U2AF1^S34F^- or SF3B1^K666N^-expressing K562 cells after pulse-labeling with BrdU. As shown in Fig. 4F and G, prolonged SMG1i treatment (3 days) increased the G2/M population in both control WT cells and spliceosome mutant cells; however, this effect was much greater in mutant cells than in WT cells. Similarly, although SMG1i treatment reduced BrdU incorporation in both WT and spliceosome mutant cells, this effect was much greater in mutant cells than in WT cells (Suppl. Fig. 4B and C). These data suggest that the combination of splicing dysregulation and NMD inhibition compromises cell cycle progression and DNA replication. To further assess the effects of NMD inhibition on DNA replication in spliceosome mutant cells, we performed DNA fiber analysis of nascent DNA after a sequential IdU/CIdU pulse-labeling procedure^69,70^. In the DNA tracts with both IdU and CldU signals, the average length of the CIdU tracts represents the overall speed of fork elongation, while the ratio of CIdU/IdU tract lengths reflects the “smoothness” of fork progression (with a ratio < 1 indicative of fork obstruction that occurs during CldU incorporation↓note that forks stalled/collapsed during the first IdU incorporation are less likely to proceed to have subsequent CIdU incorporation.)^71–73^. As shown in Fig. 4H and I, SMG1i treatment reduced the overall speed of fork progression, with a greater effect observed in spliceosome mutant cells than in control WT cells. SMG1i treatment also caused much more reduction in the CIdU/IdU ratio in spliceosome mutant cells than in control WT cells (Fig. 4J and K), indicative of replication obstruction. Defects in replication often cause fork collapse, resulting in chromosomal instability. Consistently, we detected a higher level of chromosome abnormalities, including chromosomal breaks and fusions, in K562 cells expressing U2AF1^S34F^ or SF3B1^K666N^, compared to control cells expressing WT proteins, after SMG1i treatment (Fig. 4L and M). Taken together, these data suggest that disruption of NMD in spliceosome mutant cells causes an elevated level of replication obstruction, leading to slowed replication, cell cycle arrest and chromosomal instability. These effects are likely partially responsible for the observed hypersensitivity of spliceosome mutant cells to NMD inhibition.

### R-loops are important for the hypersensitivity of spliceosome mutant cells to NMD disruption

To further define the molecular mechanisms for the sensitivity of spliceosome mutant cells to NMD disruption, we investigated the possible involvement of R-loops (a structure containing an RNA:DNA hybrid with displaced ssDNA) that are known to be elevated in spliceosome mutant cells^74–76^. Although R-loops participate in a number of physiological processes, abnormal R-loop formation can interfere with DNA replication and transcription, causing DNA damage, genomic instability and cell death^77,78^. Consistent with this notion, spliceosome mutant cells exhibit intrinsic DNA damage, cell cycle arrest and chromosomal instability (Fig. 4F-M)^79–83^. It has been shown previously that NMD factors regulate levels of telomeric repeat-containing RNA (TERRA) on telomeres, suggesting that they play a role in R-loop regulation at telomeres^84^. Notably, we observed that SMG1i treatment or UPF1 depletion resulted in a marked increase in overall R-loop levels in cells (without spliceosome mutations) (Fig. 5A and B, Suppl. Fig. 5A and C). UPF1 depletion and SMG1i treatment also caused increased H2AX phosphorylation (γH2AX), a marker of DNA damage (Suppl. Fig. 5B and D). These increased levels of both R-loops and γH2AX were largely rescued by overexpression of RNase H1 that removes R-loops and has been shown to rescue R-loop associated genomic instability (Suppl. Fig. 5E-H)^85,86^. Thus, both spliceosome mutations and NMD disruption cause R-loop accumulation and DNA damage. This raises the possibility that the combination of spliceosome mutations and NMD inhibition causes even more abnormal R-loops and DNA damage. Indeed, SMG1i treatment and U2AF1^S34F^ or SF3B1^K700E^ mutations exhibited additive effects on the levels of R-loops and γH2AX in U2OS cells (Fig. 5C, D, F, G). These effects were largely rescued by overexpression of RNase H1 (Fig. 5C, D, F, G). Remarkably, the selective killing effect of SMG1i on spliceosome mutant K562 cells expressing U2AF1^S34F^ or SF3B1^K666N^ was also largely rescued by RNase H1 overexpression, indicating that R loops are a major underlying mechanism for the sensitivity of spliceosome mutants to NMD inhibition (Fig. 5E and H). Taken together, the results described above strongly suggest that disruption of NMD in spliceosome mutant cells causes further increase in R loops, leading to replication defects, DNA damage, chromosomal instability and cell death.

**Figure 5.**
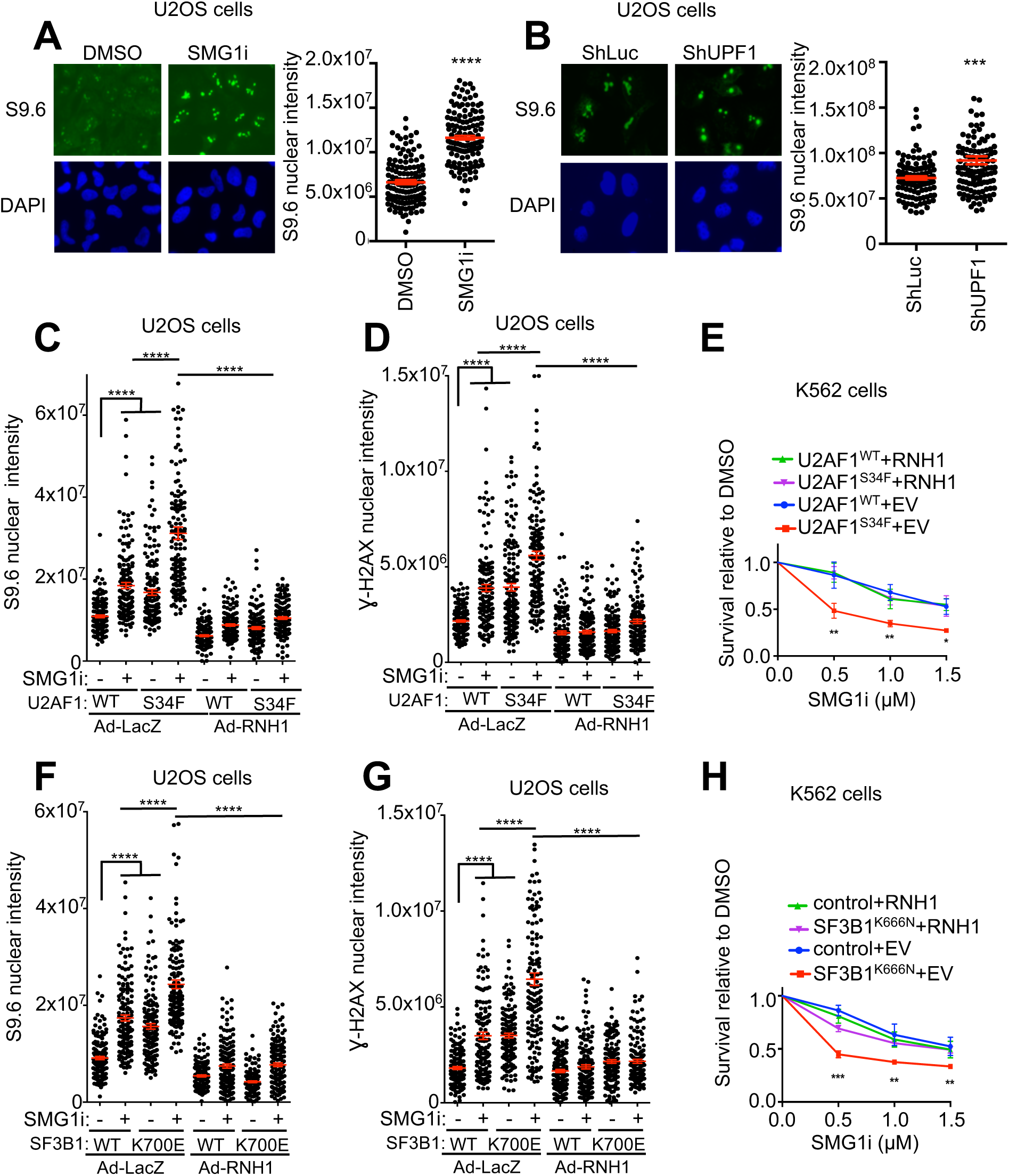
Elevated R-loop formation is a major underlying mechanism for the hypersensitivity of spliceosome mutant cells to NMD inhibition. A. Effects of SMG1i treatment on R-loops. U2OS cells were treated with SMG1i (5 μM) for 24 hours and then immunofluorescence was performed to detect nuclear S9.6 signal. Left, representative images showing nuclear signal of S9.6. Right, Quantification of nuclear S9.6 signal. Red bars represent the mean ± S.E.M of two independent experiments. A total of 130 nuclei analyzed per sample. ****p ≤ 0.0001 (Mann Whitney test). B. Effects of shRNA-mediated knockdown of UPF1 on R-loops. U2OS cells were infected with shLuc- or shUPF1-expressing lentiviruses and then incubated for 5 days. R-loops in the nucleus were detected by immunofluorescence staining using the S9.6 antibody. Left, representative images showing nuclear signal of S9.6. Right, Quantification of nuclear S9.6 signal. Red bars represent the mean ± S.E.M of two independent experiments. A total of 130 nuclei analyzed per sample. ***p ≤ 0.001 (Mann Whitney test). C. Effects of RNH1 expression on R-loops in SMG1i-treated cells expressing U2AF1^WT^ or U2AF1^S34F^. U2OS cells stably expressing U2AF1^WT^ or U2AF1^S34F^ were infected with adenovirus expressing LacZ control or RNH1 and then treated with SMG1i (1 μM) for 3 days. R-loops in the nucleus were detected by immunofluorescence staining using the S9.6 antibody. Red bars represent the mean ± S.E.M of two independent experiments. A total of 150 nuclei analyzed per sample. ****p ≤ 0.0001 (Mann Whitney test). D. Effects of SMG1i treatment on γH2AX in cells expressing U2AF1^WT^ or U2AF1^S34F^. U2OS stably expressing U2AF1^WT^ or U2AF1^S34F^ Cells were infected with adenovirus expressing lacZ control or RNH1 and then treated with SMG1i (1 μM) for 3 days. γH2AX levels were detected by immunofluorescence staining. Red bars represent the mean ± S.E.M of two independent experiments. A total of 150 nuclei analyzed per sample. ****p ≤ 0.0001 (Mann Whitney test). E. Effects of SMG1i treatment (3 days) on the viability of K562 cells stably expressing EV/RNH1 after induction of U2AF1^WT^ or U2AF1^S34F^. Data represent the mean ± SD of three independent experiments. **p ≤ 0.01; *p ≤ 0.05 (unpaired t-test). F. Effects of RNH1 expression on R-loops in SMG1i-treated cells expressing SF3B1^WT^ or SF3B1^K700E^. U2OS cells stably expressing SF3B1^WT^ or SF3B1^K700E^ were infected with adenovirus expressing lacZ control or RNH1 and then treated with SMG1i (1 μM) for 3 days. R-loops in the nucleus were detected by immunofluorescence staining using the S9.6 antibody. Red bars represent the mean ± S.E.M of two independent experiments. A total of 150 nuclei analyzed per sample. ****p ≤ 0.0001 (Mann Whitney test). G. Effects of RNH1 expression on γH2AX in SMG1i-treated cells expressing SF3B1^WT^ or SF3B1^K700E^. U2OS cells stably expressing SF3B1^WT^ or SF3B1^K700E^ were infected with adenovirus expressing lacZ control or RNH1 and then treated with SMG1i (1 μM) for 3 days. γH2AX levels were detected by immunofluorescence staining. Red bars represent the mean ± S.E.M of two independent experiments. A total of 130 nuclei analyzed per sample. ****p ≤ 0.0001 (Mann Whitney test). H. Effects of RNH1 expression on the viability of wild type or SF3B1^K666N^ knock-in K562 cells treated with SMG1i. Wild type or SF3B1^K666N^ knock-in K562 cells were infected with adenovirus expressing lacZ control or RNH1 and then treated with SMG1i for 3 days. Data represent the mean ± SD of three independent experiments. ***p ≤ 0.001; **p ≤ 0.01; *p ≤ 0.05 (unpaired t-test).

## Discussion

In this study, we have developed a new reporter system for NMD in human cells, identified many putative NMD factors and regulators, and uncovered a synthetic lethal relationship between defective general splicing and NMD disruption. Our fluorescence- and bioluminescence-based reporter system can measure NMD activity in individual cells as well as in cell populations. Using this reporter we performed a genome-wide CRISPR/Cas9 knockout screen and identified many putative NMD-promoting factors, including components of the U2 spliceosome and other factors required for early spliceosome assembly (Fig. 1, Fig. 2, Suppl. Fig.1, Suppl. Fig. 2). These hits, together with those identified previously in CRISPR/Cas9 knockout and siRNA knockdown screens using different reporter systems, provide a rich resource for future characterization of the mechanism and regulation of NMD in human cells^59,87^. Interestingly, our data indicate that NMD activity is partially attenuated in cells harboring mutations in U2 spliceosome components *SF3B1* or *U2AF1*, which are common in MDS and cancers. These spliceosome mutant cells are preferentially sensitive to NMD disruption, suggesting that NMD is a unique therapeutic vulnerability for malignancies with defects in general splicing.

The enrichment of early spliceosome factors on the hit lists of our CRISPR/Cas9-based screen and the siRNA-based screen by Hogg and colleagues sheds new light on the relationship between NMD and RNA splicing^59^. Because both introns and EJC assembly are important for NMD of multiple reporters tested, it is believed that pre-mRNA splicing plays a crucial role in NMD in mammals, at least for some transcripts. However, it remains unclear whether NMD of those transcripts requires the complete process of splicing, or whether the association of certain spliceosome factors to pre-mRNA (which then facilitates EJC assembly) during the splicing process promotes NMD. The results of two genome-wide screens with different reporter systems and gene disruption methods suggest that the latter possibility is more likely to be correct. Six out of the 9 verified top hits in our screen *(SF3B1, SF3B5, SF3A3*, *PRPF19*, *RNF113A*, *DGCR14*) are spliceosome factors required for early steps of splicing, and 3 of these belong to U2 snRNP^88,89^. The top 50 hits of potential NMD factors or regulators in the screen by Hogg and colleagues include 6 components of the U2 spliceosome, 4 PRP19-related genes, and 4 genes involved in catalytic steps of splicing/complex C^59^. It is possible that these identified splicing factors represent the proteins that promote the recruitment or maintenance of EJC required for NMD. Consistent with this idea, the core EJC factors (Magoh, RBM8A, and eIF4A3) were shown to be present in spliceosome B complex, and more stably associated with the C complex^90^. The recruitment of the core EJC factor eIF4A3, which is believed to initiate EJC assembly, is facilitated by the splicing factor CWC22 (a top hit in our screen)^91–93^. Interestingly, CWC22 has also been shown to interact with PRPF19 and Slu7 (a top hit in the Hogg screen), although it is not clear whether these interactions are important for eIF4A3 recruitment ^92,94^. The spliceosome factors identified in our two screens may also facilitate the recruitment of the other three core EJC factors (MAGOH, *RBM8A* and MLN51) to mRNA, which apparently occurs separately from the CWC22-mediated recruitment of eIF4A3. Alternatively, these spliceosome factors may promote NMD by maintaining the stability of EJC on mRNA or even through an EJC-independent mechanism. Further validation and characterization of these spliceosome factors will provide mechanistic insights into the functional interplay between RNA splicing and NMD.

While the normal RNA splicing process promotes NMD, dysregulation of general splicing can generate widespread nonsense mRNAs that rely on NMD for clearance. Interestingly, we found that NMD activity is attenuated in cells expressing SF3B1(K700E) or U2AF1(S34F), both of which are frequently found in MDS, AML, CMML and solid tumors ^26,95–97^. Our results are in agreement with a previous observation that the levels of certain NMD factor transcripts (which are themselves NMD targets) were increased in cells expressing U2AF1(S34F) mutant^9,98^. Spliceosome mutations cause largely distinct patterns of alterations in splicing and gene expression, but a shared feature is the generation of numerous aberrant nonsense mRNAs^9,17,18^. The attenuated NMD activity observed in cells expressing mutant spliceosome factors may be caused by the loss of ability of these factors to promote NMD. Alternatively, abnormal splicing/expression of NMD factors (e.g., UPF3A and SMG7^99^) and/or saturation of the NMD machinery by the high levels of nonsense mRNAs in these mutant cells may lead to decreased NMD efficiency. However, certain splicing factor mutations can also enhance NMD of specific mRNAs. A recent study by Krainer and colleagues shows that SRSF2^P95H/L/R^, which are also found in MDS and cancers, stimulate NMD of target transcripts that contain binding sites of these mutants ^100^.

The prevalence of nonsense mRNAs in spliceosome mutant cells and the requirement of NMD for their clearance raise the possibility that functional disruption of NMD can selectively kill these cells. Indeed, we found that cells expressing SF3B1^K700E^, SF3B1^K666N^ or U2AF1^S34F^ were much more sensitive to UPF1 knockdown or SMG1 inhibition, compared with cells expressing WT proteins (Fig. 4A-E). This preferential sensitivity is correlated with the phenotypes of cell cycle arrest, DNA replication defects, DNA damage and chromosomal instability observed in these mutant cells (Fig. 4F-M). Remarkably, the sensitivity of spliceosome mutant cells to SMG1 inhibition could be rescued by RNaseH1 overexpression (Figs. 5E and H), suggesting that R loops are a major underlying mechanism of the synthetic lethality between defective splicing and NMD disruption. In support of this idea, we found that SMG1 inhibition and spliceosome mutations in combination caused an additive effect on R loop formation (Fig. 5A-C, F, Suppl. Fig.5A and C)^76,82,83^. This effect on R loops is in line with that on γH2AX, a marker of DNA damage that is induced by abnormal R loops (Fig. 5D and 5G). Thus, our data strongly suggest that inhibition of NMD in spliceosome mutant cells causes a further increase in R loop levels, which in turn causes heightened DNA replication obstruction, DNA damage, chromosomal instability and cell death.

The selective killing effects of NMD inhibition on spliceosome mutant cells suggest that NMD is an attractive target for treating certain MDS and cancers. Based on the observation that spliceosome mutant cells are generally more sensitive to further splicing perturbation, a major effort has been focused on developing splicing modulators such as E7107 and H3B-8800↓both of which bind to SF3B1↓as therapies for MDS, AML and CMML^101–105^. However, clinical trials with these compounds either were suspended due to toxicity, or did not achieve objective responses^102–105^. Our results suggest the possibility that NMD inhibition, more specifically SMG1 inhibition, is an alternative strategy for treating MDS and cancers with defective splicing (Fig. 4C-E). It is worth noting that although complete disruption of NMD appears to be lethal, its attenuation is tolerated and occurs normally in certain developmental processes and in the responses to cellular stress^33,58,106,107^. Beyond the cell-autonomous effects described in this study, NMD inhibition also has the potential to induce anti-cancer immunity by increasing the production of cancer neoantigens encoded by mis-spliced nonsense mRNAs in spliceosome mutant cells^108–110^. In addition to splicing modulation and NMD inhibition, spliceosome mutant cells are also sensitive to inhibition of ATR, another SMG1-related kinase in the PIKK family^83,111,112^. Of note, SMG1i described in this study is highly specific for SMG1 (Suppl. Fig 3B). At the concentrations used for NMD inhibition in this study, SMG1i does not inhibit ATR (Suppl. Fig. 3C). It will be important to directly compare different therapeutic strategies for MDS and cancers with defective splicing and test the potential of combination treatment with spliceosome modulators, SMG1i and ATRi.

## Acknowledgments

We thank Won Kyun Koh for his contribution in generating the new NMD reporter containing both fluorescent proteins and luciferases. We are grateful to Dr. Niels H. Gehring for providing the 4boxB and λN-V5-UPF3B expression constructs, and to Dr. Sheila Stewart for providing recombinant adenoviruses expressing RNH1. This work was supported by an NIH grant (R01GM098535) and Siteman Investment Program Awards (4036, 5124) from Washington University to Z.Y., a Developmental Research Program (DRP-1901) of the SPORE in Leukemia (NIH/NCI, P50CA171963) and the Edward P. Evans Foundation to M.W. and Z.Y. Abigael Cheruiyot was a Howard Hughes Medical Institute International Student Research fellow from 2016-2019. SMG1i was provided by Amgen, Inc. We thank Drs. Rick Austin and Tim Cushing for leading efforts at Amgen to develop the SMG1 inhibitor used in this study. Support for procurement of human samples was provided by the Genomics of AML Program Project of the National Cancer Institute (P01 CA101937).

## Author Contributions

Z.Y. conceived and supervised the overall project. A.C. and S. L. conducted all the experiments and analyzed the results with contributions from S.N.S., T.A., D.S.L.,Y.L. Z.Y., B.A.W., and W.W. under the supervision of Z.Y., M.J.W., F.X., D.H. and D.C.L. Y.C. and X.W. performed bioinformatic analysis of the CRISPR/Cas9 knockout screen results together with A.C. S.P-M. and E.A.O. generated genetically modified K562 SF3B1^K666N^ mutant cells. M.J.W., X.W., J.M.B., F.X., D.H., E.A.O. and D.C.L. provided key reagents and/or critical discussions on the project. A.C. and Z.Y. wrote the manuscript with input from all the other authors. All authors read and approved the manuscript.

## Materials and Methods

### Key reagents, oligo sequences, genetically modified K562 SF3B1^K666N^ cells, SMG1 inhibitor

The key reagents used in this study are listed in Table 1. The sequences of sgRNAs, RT-qPCR primers and *oligos for CRISPR/Cas9 knock-in* are listed in Table 2. Genetically modified K562 *SF3B1^K666N^* cells were generated using CRISPR-Cas9 technology. Briefly, 200,000-400,000 K562 cells were transiently co-transfected with 150 ng of sgRNA (Synthego), 75 ng Cas9 protein (Berkeley Macrolab), 100 pmol of ssODN donor (hSF3B1.K666N.anti.ssODN), and 200 ng pMaxGFP expression plasmid (Lonza, #13429329) via nucleofection (Lonza, 4D-NucleofectorTM X-unit) using solution P3 (Lonza, #V4XP-3024), program FF-120 in small cuvettes according to the manufacturer’s recommended protocol. To obtain heterozygous clones, two ssODN donors were used: one donor containing the desired modification and a blocking modification to prevent subsequent editing after incorporation of the desired modification, and another donor containing only a silent blocking modification in order to prevent editing by non-homologous end joining on the second allele. Five days post nucleofection, cells were single-cell sorted by FACS to enrich for GFP+ (transfected) cells, clonally selected, and verified for the desired targeted modification via targeted deep sequencing and analysis with *CRIS.py*^113^. The SMG1i used in this study is an ATP-competitive sulfonamide compound developed by Amgen, Inc.

### Cell culture, transfection, lentivirus production and infection

Human cell lines were cultured in DMEM (Sigma, D5796) (for U2OS, HEK293T and Calu-6) or in RPMI 1640 medium (Gibco, 1187) (for K562) supplemented with 10% fetal bovine serum (FBS), 100 units/ml penicillin, 100 µg/ml streptomycin in a 5% CO2 incubator at 37 °C.

Human GeCKOv2 CRISPR knockout pooled library (Addgene # 1000000049) and lenti-Cas9 plasmid were obtained from addgene (Addgene # 52962), a gift from Dr. Feng Zhang^114,115^. The library was amplified in DH5α cells on 100 agar plates (15 cm). Deep Sequencing results indicate 98% coverage of all the sgRNAs in the designed library. To generate lentiviruses expressing shRNA, Cas9, sgRNA-Cas9, or the GeCKOv2 sgRNA library, HEK 293T cells were transfected with lentiviral vectors and packaging vectors (pCMV-VSVG and psPAX2, Addgene # 8454 and 12260, respectively), using the Mirus TransIT-LT1 transfection reagent. Cell culture medium containing lentiviruses was collected 48 and 72h after transfection, filtered (0.45 μm filter, Millipore Sigma #SLHV033RS) and used for infection of target cells in the presence of 8 μg/mL polybrene.

Transfection of sgRNA-Cas9 plasmids into Calu-6 cells for individual gene validation was done using Lipofectamine 3000 transfection reagent (ThermoFisher Scientific), according to manufacturer’s protocol. Cells were selected with puromycin (3 μg/mL) 48 hours after transfection. RNA stability was analyzed 6 days after transfection via an actinomycin D chase assay, as described previously^35,49^.

### Generation of a fluorescence/bioluminescence-based NMD reporter

The fluorescence/bioluminescence-based NMD reporter system used for the genome-wide CRISPR/Cas9 screen in this study was generated by inserting ORFs of mCherry and EGFP immediately upstream of CBR and CBG, respectively, into our previously described bioluminescence-based NMD reporter^35,48^. Complete sequence and annotation of the reporter are available upon request. U2OS cells stably expressing this NMD reporter system was generated by co-transfecting the NMD reporter plasmid and a pMXs-puro vector that encodes a puromycin-resistance gene into U2OS cells, using Mirus TransIT-LT1 transfection reagent. After selection with puromycin (1.5 μg/mL), single clones expressing the reporter were isolated and validated by examining the effects of depletion of known NMD factors or caffeine treatment on NMD of the integrated reporter.

### Assays for NMD of the fluorescence/bioluminescence-based reporter

Multiple assays were used to measure NMD activity using our new fluorescence/ bioluminescence-based reporter. For live cell imaging, mCherry and EGFP fluorescence signals in U2OS reporter cells plated in 3.5 cm glass-bottomed dishes (MatTek corporation) were acquired using a Nikon Eclipse TiE inverted microscope with MetaMorph software, as described previously^116^. For flow cytometry, resuspended single U2OS reporter cells were analyzed on a FACS machine (Sony, Synergy HAPS 1) to separate cell populations based on mCherry and EGFP signals. For western blots, anti-HA antibodies were used to detect reporter fusion proteins HA-mCherry-CBR-TCR(PTC) and HA-EGFP-CBG-TCR(WT). For RT-qPCR, total RNA was isolated using RNAqueous™ Total RNA Isolation Kit from ThermoFisher Scientific (AM1912), or TRIzol™ reagent from ThermoFisher Scientific (15596). Trace DNA contamination was removed using TURBO DNA-free™ Kit from ThermoFisher Scientific (AM1907), followed by reverse transcription to synthesize cDNA using PrimeScript RT kit from Clontech (RR037A). qPCR was performed using a two-step PCR protocol (melting temperature: 95°C; annealing/extension temperature: 60°C; cycle number: 40) on an ABI V117 real-time PCR system with PowerUp SYBR Green Master Mix (ThermoFisher Scientific, A25742). The mRNA levels of the housekeeping gene GAPDH was used for normalization. The sequences of the primers used are listed in Table 2.

### Knockdown of NMD factors

To knock down NMD factors in U2OS reporter cells for reporter validation, lentiviruses expressing previously validated shRNAs targeting UPF1 or SMG1 were used^35^. A shRNA targeting firefly luciferase (shLuc) was used as a control. The sequences of shRNA used are listed in Table 2. U2OS reporter cells were infected with shRNA and incubated for 4 days before NMD reporter analysis. Depletion of the proteins was confirmed via western blot analysis using anti-SMG1 (Cell Signaling Technology, 9149), and anti-UPF1 (Cell Signaling Technology, 12040) antibodies. Tubulin (Santa Cruz, sc8035) was used as a loading control.

### Pooled genome-wide CRISPR/Cas9 knockout screen to identify new NMD factors and regulators

U2OS reporter cells were infected with lentiviruses expressing Cas9, and then selected with blasticidin (10 μg/mL) for 5 days to establish the U2OS reporter Cas9 cell line. This cell line was validated by examining the effects of sgRNAs targeting SMG1 or UPF2 on NMD of the reporter. To carry out the screen, U2OS reporter Cas9 cells were infected with lentiviruses expressing the GeCKOv2 library (containing two sub-libraries) at a MOI of less than 1 with a 500x coverage of the library. Six days after infection, FACS was performed to collect cells with increased mCherry to EGFP ratio, indicative of NMD inhibition. Genomic DNA was extracted using PureLink Genomic DNA kit (ThermoFisher Scientific, K182001) followed by two PCR reactions to prepare samples for Illumina Next-Gen sequencing. The first PCR was used to amplify out sgRNA inserts in the cells, and the second PCR was used to add Illumina sequencing tags as well as indexes for sample identification. To improve complexity of the library required for deep sequencing, a mixture of 5 forward primers with staggered nucleotides immediately upstream of the sgRNA sequences was used in the second PCR. The sequences of primers used are listed on Table 2. All PCR reactions were performed using Phusion Hot Start II High-Fidelity DNA Polymerase (ThermoFisher Scientific, F549L). PCR samples were sequenced using the Illumina HiSeq 2500 platform. As a baseline control for the abundance of sgRNAs in the collected cells, a fraction of unsorted cells were subjected to the same procedure of genomic DNA isolation, PCR and Next-Gen Sequencing.

### Screen data analysis and hit identification and validation

A custom Perl script was used to determine the read counts for each sgRNA and map the sgRNAs to their gene IDs in the reference GeCKOv2 library. The script is available upon request. Only the sgRNAs with at least 10 reads in each sample was used for further analysis. Analysis of genes enriched in FACS-collected cells (with inhibited NMD) compared to baseline control was performed using MAGeCK, a computational tool designed to rank genes based on the enrichment of individual sgRNAs as well as the number of enriched sgRNAs for each gene^117^.

To validate genes identified in the screen, two gRNAs from a different human gRNA library (AVANA library)^118^ for each of the 9 genes were cloned into the pLentiCRISPR V2 vector that also expresses Cas9 (Addgene, #52961). Two non-targeting gRNAs were also cloned into the same vector to serve as controls. All the sequences for the gRNAs used are listed in Table 2. Lentiviruses expressing each gRNA (and Cas9) were generated and transduced into U2OS reporter cells to deplete expression of individual genes. NMD activity was then analyzed by western blot to determine the ratio of HA-mCherry-CBR-TCR(PTC) to HA-EGFP-CBG-TCR(WT). Since the ratio was similar for both non-targeting controls, the relative NMD activity was normalized to one non-target control (sgNT-1). As an independent NMD analysis system for further validation, human pulmonary adenocarcinoma Calu-6 cells that express a PTC-containing p53 mRNA was used^35,41,49,56^. The same sgRNAs-Cas9 in pLentiCRISPR V2 described above were transfected into Calu-6 cells, followed by 48 hours of puromycin selection, to deplete expression of the top ranked 9 genes individually. Six days after transfection, cells were treated with actinomycin D (5 μg/mL) for 6 hours to inhibit transcription. RNA samples were collected immediately before and after actinomycin D treatment. RT-qPCR was performed for p53 mRNA to measure the percent mRNA remaining after addition of actinomycin D. In addition to mutant p53 mRNA, the RNA stability of several known physiological NMD targets, including *ATF4, PIM3*, *UPP1*, as well as ORCL (a control that is not a NMD target), were also measured in the same samples^57^.

### λN-boxB tethering reporter and NMD analysis

The EGFP-boxB reporter construct was generated based on the β-globin-boxB tethering reporter developed by Dr. Niels Gehring and colleagues^61^. The 3’UTR containing four boxB sites were cloned into pEGFP-C1 vector to replace the 3’ UTR of the EGFP transcription cassette. The resulting EGFP-boxB expression cassette was then sub-cloned into the pCDH lentiviral vector at Xba I and Sal I sites. A construct with scrambled boxB (boxB’) sequences was also generated as a control. To generate a Dox-inducible λN-UPF3B expression construct, the λN-V5-UPF3B ORF in pCl-neomycin-λN-V5-UPF3B (also a gift from Dr. Gehring) was PCR amplified and inserted into the pCW lentiviral vector. pCW-λN-V5 expressing λN alone, and pCW-UPF3B, without λN, were also generated as controls. Inducible U2OS cell lines expressing λN, λN-UPF3B or λN-unfused UPF3B were generated by lentivirus infection. Stable expression of EGFP-boxB or EGFP-boxB’ reporters in these cells lines were generated by lentiviral infection. Expression of the inducible proteins was induced by 1 μg/mL doxycycline for 48 hours. The stability of EGFP-boxB or EGFP-boxB’ reporter transcripts was determined by RT-qPCR analysis of RNA samples before or after 1-2 hours of actinomycin D treatment. To determine whether expression of SF3B1/U2AF1 mutants affected the decay of UPF3B-tethered RNA, cells were infected with lentivirus expressing Flag-tagged SF3B1^WT^, SF3B1^K700E^, U2AF1^WT^, or U2AF1^S34F^ 5 days prior to doxycycline induction and RNA stability analysis.

### Cell viability analysis

U2OS cells stably expressing Flag-SF3B1^WT^ or Flag-SF3B1^K700E^ were generated by lentiviral infection. To assess the sensitivity of these cells to SMG1i or PB, cells were plated in triplicates at similar density and 24 hours later treated with DMSO, SMG1i or PB. K562 cells with inducible expression of U2AF1^WT^ or U2AF1^S34F^ were generated previously^119^. To assess the sensitivity of these cells to SMG1i or PB, cells were grown in T25 flasks in the presence or absence of 250 ng/mL doxycycline for 48 hours. An equal number of induced or uninduced cells were then plated and treated with SMG1i, PB, or DMSO. K562 cells with SF3B1^K666N^ knock-in mutation were generated using the CRISPR/Cas9 technique (see above). The sensitivity of these cells and their parental K562 cells to SMG1i or PB was also evaluated. In all these experiments, alamar blue assay was performed 3 days after addition of SMG1i or PB. Relative viability of SMG1i- or PB-treated cells were obtained after normalization to DMSO-treated cells. To determine the effect of RNase H1 over-expression on cell sensitivity to SMG1i, K562 cells with inducible expression of U2AF1^WT^ or U2AF1^S34F^, or with SF3B1^K666N^ knock-in mutation were infected with lentiviruses generated from empty pCDH vector (EV), or pCDH-RNase H1 (RNH1). Cell viability in EV-, or RNH1-expressing cells after SMG1i treatment was then measured as described above.

To determine the sensitivity of spliceosome mutant cells to UPF1 depletion, U2OS cells stably expressing SF3B1^WT^, SF3B1^K700E^, U2AF1^WT^, or U2AF1^S34F^ were infected with UPF1 shRNA, or with shLuc control. Twenty hours after infection, an equal number of cells were plated and alamar-Blue assay was performed on day 3 to day 6 post infection. To determine cell growth from day 3 to day 6, the values obtained from alamar blue assay were normalized to day 3 for each cell line. To determine the relative cell viability of cells expressing WT versus mutant splicing factors after UPF1 depletion, values from cells infected with shUPF1 were normalized to values from shLuc infected cells.

### Immunofluorescence to detect R-loops and γH2AX

U2OS cells were treated with DMSO or SMG1i (5 μM) for 24 hours, or infected with shUPF1/shLuc, and incubated for 5 days. U2OS cells stably expressing SF3B1^WT^, SF3B1^K700E^, U2AF1^WT^, or U2AF1^S34F^ were infected with LacZ control, or RNase H1 adenovirus and then treated with SMG1i (1 μM) for 3 days. To detect R-loops and γH2AX by immunofluorescence staining, cells plated on cover-slips were first permeabilized with PBS containing 0.2% Triton, washed with PBS and fixed with 4% PFA, and then blocked with 10% goat serum in PBS. Cells were then incubated with antibodies against S9.6 (1:100, Millipore Sigma, MABE1095) and γH2AX (1:500, Cell Signaling Technology, 9718) for 2 hours at room temperature. Cells were washed with PBS containing 0.1% Triton, and then incubated with Alexa Fluor 488-conjugated goat anti-mouse antibody (1:500, ThermoFisher, A-11001) and Alexa Fluor 568-conjugated goat anti-rabbit (1:500, ThermoFisher, A-11011) for 1 hour. Cells were counter-stained with Hoechst (ThermoFisher, H3570). Images were acquired using Nikon Ti-E fluorescence microscope and Metamorph software (Molecular Devices). Nuclear signal of S9.6 and γH2AX were quantified using ImageJ software.

### Detection of R-loops in U2OS cells using slot blot analysis

U2OS cells were treated with DMSO or SMG1i (5 μM) for 24 hours, or infected with shUPF1/shLuc and incubated for 5 days, followed by genomic DNA extraction using phenol/chloroform/isoamyl alcohol. Genomic DNA (2 μg) was then treated with buffer alone or with 1 unit RNase H enzyme (NEB, MO297S) for 2 hours at 37 °C, and 0.2-1 μg of treated genomic DNA was then slotted directly onto 2 nylon membranes. One membrane, used as loading control was incubated in DNA-denaturing buffer (0.5 M NaOH, 1.5 M NaCl) for 10 min and neutralized for 10 min in neutralizing buffer (0.5 M Tris-HCl, 1.5 M NaCl, pH 7.2). Both membranes were UV-crosslinked, blocked with casein buffer, and then incubated with S9.6 antibody to detect R-loops, or with anti-ssDNA antibody (Millipore Sigma, MAB3868) to detect total DNA. Images were obtained using ChemiDoc imaging system, and the S9.6/ssDNA signal was quantified using ImageJ software. To determine if RNase H1 can remove R-loops in cells, U2OS cells were infected with adenovirus expressing LacZ control or RNase H1. After UPF1 knockdown or SMG1i treatment, genomic DNA was extracted and slot blot analysis was performed as described above.

### Immunoblotting

Detection of all the proteins was done using SDS-PAGE and immunoblotting using the Odyssey Infra-red Imaging System as described before^120^. Briefly, cells were lysed using 50 mM Tris, 10% glycerol, 2% SDS, 5% β-mercaptoethanol. Protein lysates were run on SDS-PAGE gels and transferred to a PVDF membrane, blocked with casein buffer and incubated with the indicated primary antibodies (see Table 1). DyLight 800- and DyLight 680-conjugated secondary antibodies were used to detect the primary antibodies, and signals were acquired using Odyssey Infrared Imaging System (Li-COR Biosciences).

### BrdU incorporation and cell cycle analysis

Cell cycle and DNA replication analysis was performed by flow cytometry after BrdU incorporation, as described before^69^. K562 cells expressing inducible U2AF1^WT^ or U2AF1^S34F^, or with SF3B1^K666N^ knock-in mutation were treated with 1 μM SMG1i for 3 days, and then incubated with BrdU (20 μM) for 30 min. Subsequently, cells were washed with cold PBS and fixed with 70% ethanol overnight. DNA was denatured using 2 N HCl/0.5% Triton X-100 for 30 minutes in room temperature, and then neutralized with 0.1 M sodium tetraborate, pH 8.5. Cells were then incubated overnight with mouse anti-BrdU antibody (1:00, BD Biosciences, 347580) in PBS + 0.5% Tween 20 + 1% BSA. After washing with PBS + 1% BSA, cells were incubated with Alexa Fluor 488-conjugated goat anti-mouse IgG (1:500, ThermoFisher, A-11001) for 1 h. After washing with PBS + 1% BSA, cells were re-suspended in PBS containing propidium iodide (20 μg/ml) and RNase A (200 μg/ml) and incubated at 37°C for 30 min. Flow cytometry was performed using BD FACSCalibur Flow Cytometer and cell cycle profile was analyzed with FlowJo software.

### Metaphase chromosome spreads

Metaphase chromosome spreads to evaluate genomic instability following SMG1i treatment were performed as previously described^70^. K562 cells expressing inducible U2AF1^WT^ or U2AF1^S34F^, or SF3B1^K666N^ knock-in mutation were treated with 1 μM SMG1i for 2 days. Cells were then washed and incubated in fresh medium for another 2 days. In the last 4 hours, cells were incubated with 10 μM nocodazole. Cells were then collected, washed with PBS, and re-suspended with10 ml of pre-warmed hypotonic solution (10 mM KCl, 10% FBS) and incubated for 10 min at 37 °C. Cells were fixed by adding 500 μl of cold fixation buffer (1V acetic acid: 3V methanol), and washed in the same buffer 4 times before overnight incubation in the same buffer at 4 °C. The nuclei were then spread on pre-chilled slides, air-dried overnight, and mounted with Prolong Gold Antifade reagent (ThermoFisher, P36930) containing Hoechst (ThermoFisher, H3570). Images were acquired using Nikon Ti-E fluorescence microscope. Fifty randomly selected metaphases per experiment (total of 3 experiments) were scored for chromosomal aberrations. Statistical analysis was done using GraphPad Prism 6.

### DNA fiber assay

DNA fiber assay to determine replication fork speed was performed as previously described^70^. K562 cells expressing inducible U2AF1^WT^ or U2AF1^S34F^, or SF3B1^K666N^ knock-in mutation were treated with 1 μM SMG1i for 3 days. Cells were then incubated with 20 μM 5-iodo-2′-deoxyuridine (IdU, Sigma-Aldrich) for 30 minutes, followed by incubation with 400 μM 5-chloro-2′-deoxyuridine (CldU, Sigma-Aldrich) for 30 minutes. Cells were then collected and washed with cold PBS, and then re-suspended at a concentration of 4×10^6^ cells/ml. A total of 2 ul of the cell solution was combined with 8 ul of lysis buffer (200 mM Tris.HCl pH 7.5; 50 mM EDTA; 0.5 % SDS) on a glass slide. Cells were allowed to settle on the slide for 5 minutes and then tilted at 20-45° angle to allow DNA to slowly spread on the slide. The resulting DNA spreads were air-dried, fixed in 3:1 methanol/acetic acid and stored at 4 °C. DNA fibers were denatured using 2.5 N HCl for 1 hr, washed with PBS and blocked with 5% BSA in PBS-T (PBS + 0.1% Tween-20) for 1 hr. CldU and IdU tracks were detected with rat anti-BrdU antibody (1:50, Abcam, ab6326), and mouse anti-BrdU antibody (1:50, BD Biosciences, 347580), respectively. Secondary antibodies (anti-rat Alexa 488, (1:100, ThermoFisher, A-11077) and anti-mouse Alexa 546 (1:100, Thermofisher, A21123) were used to detect CldU and IdU. Antibody incubations was performed in a humid 37°C chamber for 1 hr for primary antibodies, and 45 min for secondary antibodies. The slides were air-dried and mounted with Prolong Gold Antifade reagent (ThermoFisher, P36930). Fluorescence images of IdU and CldU tracks were captured using an inverted microscope (Nikon Ti-E) and Metamorph software (Molecular Devices). The IdU and CldU tract lengths were measured using ImageJ software, and the values were converted into micrometers using the scale bars created by the microscope. Fork speed was measured as the length of CldU tract in the tracts containing both IdU and CldU. Statistical analysis was performed using Prism 6.

### RNA-sequencing

K562 cells with inducible expression of U2AF1^WT^ or U2AF1^S34F^ were incubated with or without 250 ng/ml doxycycline for 48 hours. Cells were then counted and equal numbers of cells were plated and treated with DMSO or SMG1i (1μM) for 24 hours. Doxycycline was maintained during the treatment. Total RNA was extracted using Trizol LS (ThermoFisher, 10296028) and RNeasy mini kit (Qiagen, 74104). Genomic DNA was removed using Turbo DNA-Free Kit (ThermoFisher, AM1907). Total RNA integrity was determined using Agilent bioanalyzer (2100 Bioanalyzer system) using RNA 6000 Nano Kit (Agilent, 5067-1511). Library preparation was performed using 500 ng of total RNA. Ribosomal RNA was removed using Ribozero followed by RNA fragmentation, cDNA preparation and generation of stranded libraries using the TruSeq Stranded Total RNA kit with Ribo-Zero Gold, set A, RS-122-2301 (Illumina) according to the manufacturer protocol. Sequencing was performed on the NovaSeq 6000 platform (Illumina) to generate 2 × 150 bp paired-end reads. For patient samples, total RNA was extracted from unfractionated bulk bone marrow cells from four acute myeloid leukemia (AML) samples harboring a U2AF1(S34F) mutation with 62-73% myeloblasts (Patient IDs are PPI 009, UPN 245450, UPN 445045 and UPN 633734). RNA was also isolated from CD34^+^ cells that were flow-sorted from bone marrow aspirate samples of five normal healthy donors. Samples were obtained as part of studies that were approved by the Human Research Protection Office at Washington University School of Medicine and all individuals provided written informed consent. Bone marrow aspirate samples from normal healthy donors were processed via ammonium–chloride–potassium red cell lysis, washed once in phosphate-buffered saline, and then stained for flow cytometry using the CD34-phycoerythrin (PE) antibody (PE-pool, Beckman Coulter, IM1459U, Brea, CA). RNA-sequencing libraries were then prepared from RNA isolated from the bulk mutant AML bone marrow cells and the normal CD34^+^ bone marrow cells using TruSeq Stranded Total RNA kit (Illumina) described above. Sequencing was performed on the NovaSeq 6000 platform (Illumina) to generate 2 × 150 bp paired-end reads. RNA sequencing results comparing MDS patient bone marrow samples harboring SF3B1(K700E) to healthy control individuals were obtained from Pellagatti et al^121^. CD34^+^ cell isolation, RNA extraction, library preparation, and RNA sequencing for these samples was previously described^121^.

### RNAseq data analysis of K562 cells treated with SMG1i

RNAseq read quality was assessed using FastQC^122^. Reads were mapped using HISAT2 (version 2.1.0)^123^ against the GRCh37.67 version of the Human genome from Ensembl consortium. Gene level counts were processed excluding any secondary or unmapped reads using the Samtools and filtering the bam files with 0×100 flag. Resulting reads were further processed using HTseq (version 0.11.0)^124^. Differentially expressed genes were identified using DESeq2, after filtering for genes that had 10 reads in at least half the samples^125^. Genes that are significantly differentially expressed were filtered using a False Discovery rate(FDR) cut-off of <10%. Transcripts Per Kilobase Million (TPM’s) were calculated using StringTie (version 1.3.3)^126^. Gene enrichment analysis against Gene Ontology (GO) knowledgebase was performed using Clusterprofiler version (3.10.1)^127^. Pathway enrichment analysis was performed using GSEA pre-ranked method that is available with GSEA (version 4.0.3)^128,129^ against Reactome and MSIGDB gene sets.

### GSEA of endogenous NMD targets in splicing mutant cells

As described above, we performed pathway enrichment analysis against Reactome and MSIGDB gene sets. To further investigate the effect of splicing factor mutants on NMD target mRNAs, we created a custom list of gene-set (endogenous NMD targets) curated from the genes upregulated by the knockdown of the key NMD target gene UPF1^65^. Gene expression data for each target gene was used to create custom gene-set by focusing on genes that were upregulated at FDR 5% and fold-change of 2. We also created a custom gene-set for NMD target mRNAs based on the effects of SMG1 inhibition (SMG1i). To do this, we identified genes that had different expression levels between K562 cells treated with SMG1i (WT_NO_DOX_SMG1i) compared to vehicle treatment (WT_NO_DOX). Genes that were upregulated in K562(WT_NO_DOX_SMG1i) *(FDR*≤*10%, Log2FC*≥*2)* were used to create a custom gene-set.

### Identification and characterization of SMG1i

The SMG1i was identified through structure-activity relationship analysis of sulfonamide compounds determined to inhibit SMG1 enzymatic activity. The compound was generated by adding 3.0 ml trimethyl orthoacetate to a suspension of 1.9 g (2.3 mM) 3,4-diamino-*N*-(2-chloro-5-(7-cyanoimidazo[1,2-*a*]pyridin-3-yl)phenyl)benzenesulfonamide in 15 ml acetic acid. This reaction mixture was heated (120 °C for 30 minutes), cooled to ambient temperature, then concentrated. Flash column chromatography on silica gel was used to purify the final SMG1i material (*N*-(2-chloro-5-(7-cyanoimidazo[1,2-*a*]pyridin-3-yl)phenyl)-2-methyl-1H-benzo[d]imidazole-5-sulfonamide). Biochemical assays were used to evaluate the activity and specificity of the SMG1i. Inhibition of SMG1 protein kinase activity was assessed using a Homogeneous Time-Resolved Fluorescence (HTRF) assay (Cisbio). 0.8 nM Flag-tagged full-length SMG1 enzyme (produced in HEK293 cells) was incubated with 50 mM recombinantly produced avidin-labeled p53 protein (amino acids 1-83; produced in E. coli) as substrate, a concentration range of SMG1i and 30 μM ATP in reaction buffer (50 mM HEPES pH 7.5, 1 mM EGTA, 10 mM MgCl_2_, 0.01% Brij™-35 (Thermo Fisher), 1 mM DTT) in a 384-well microplate (Greiner) for 1 hour at room temperature. SMG1-mediated phosphorylation of p53, and inhibition by SMG1i, was detected by addition of 1 nM Europium-labeled anti-p53 phosphoserine 15 antibody (New England Biolabs/ Cisbio) and 40 nM streptavidin-conjugated APC (Prozyme) in Lance detection buffer (Perkin Elmer). The time-resolved fluorescence resonance energy transfer (TR-FRET) was measured on an EnVision plate reader (Perkin Elmer). Activity of the SMG1i against other PI3 kinase family members (PI3Kα, PI3Kβ, PI3Kγ, PI3Kδ, MTOR) was evaluated using Amplified Luminescent Proximity Homogeneous Assay (Alpha) Screen assays (Perkin Elmer and Echelon Biosciences). PI3K or mTOR protein (produced in Sf9 cells) was incubated with phosphatidylinositol 4,5-bisphosphate substrate with 4-10 μM ATP and a concentration range of SMG1i in Tris kinase buffer (50 mM Tris pH 7, 14 mM MgCl_2_, 2 mM Na cholate, 100 mM NaCl, 2 mM DTT) for 1.5 hours at room temperature protected from light. The phosphorylated product, phosphatidylinositol (3,4,5)-triphosphate, competes with biotinylated inositol 1,3,4,5-tetrakisphosphate on donor beads for interaction with a phosphatidylinositol (3,4,5)-triphosphate detector protein on acceptor beads. Interaction of donor and acceptor beads generates a luminescent signal that is measured using an EnVision plate reader (Perkin Elmer) with 680 nm excitation and 520-620 nm emission.

---

**Supplementary Figure 1.**
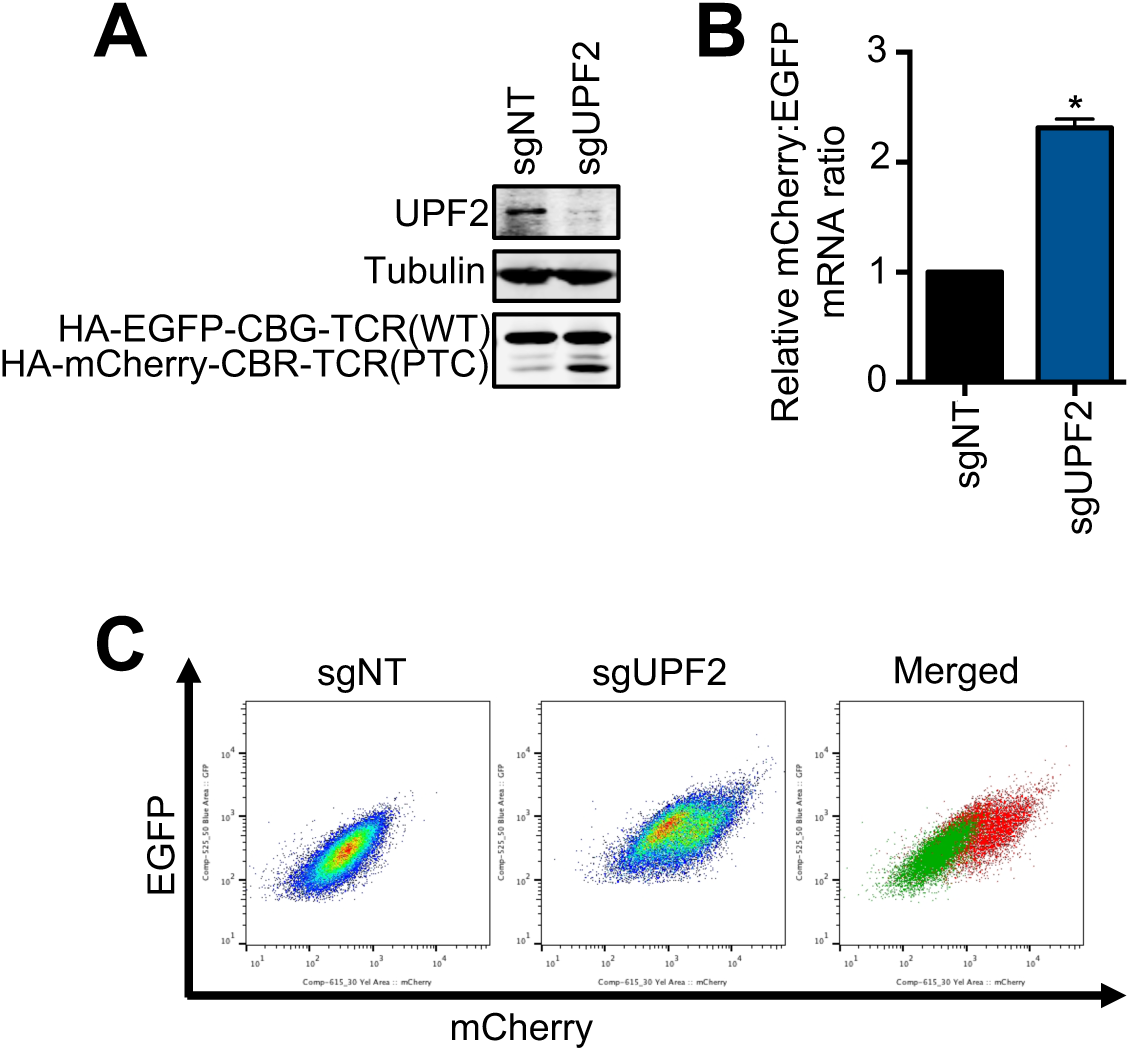
A. Western blot analysis of the protein products of the NMD reporter in Cas9-expressing U2OS reporter cells after sgRNA-mediated depletion of UPF2. B. Ratios of mCherry-containing reporter mRNA to EGFP-containing reporter mRNA in Cas9-expressing U2OS reporter cells after UPF2 depletion. The mCherry/EGFP mRNA ratio of the sgNT control was normalized to 1. Data represent the mean ± SD of three independent experiments. *p ≤ 0.05 (paired t-test). C. FACS analysis of Cas9-expressing U2OS reporter cells after UPF2 depletion.

**Supplementary Figure 2.**
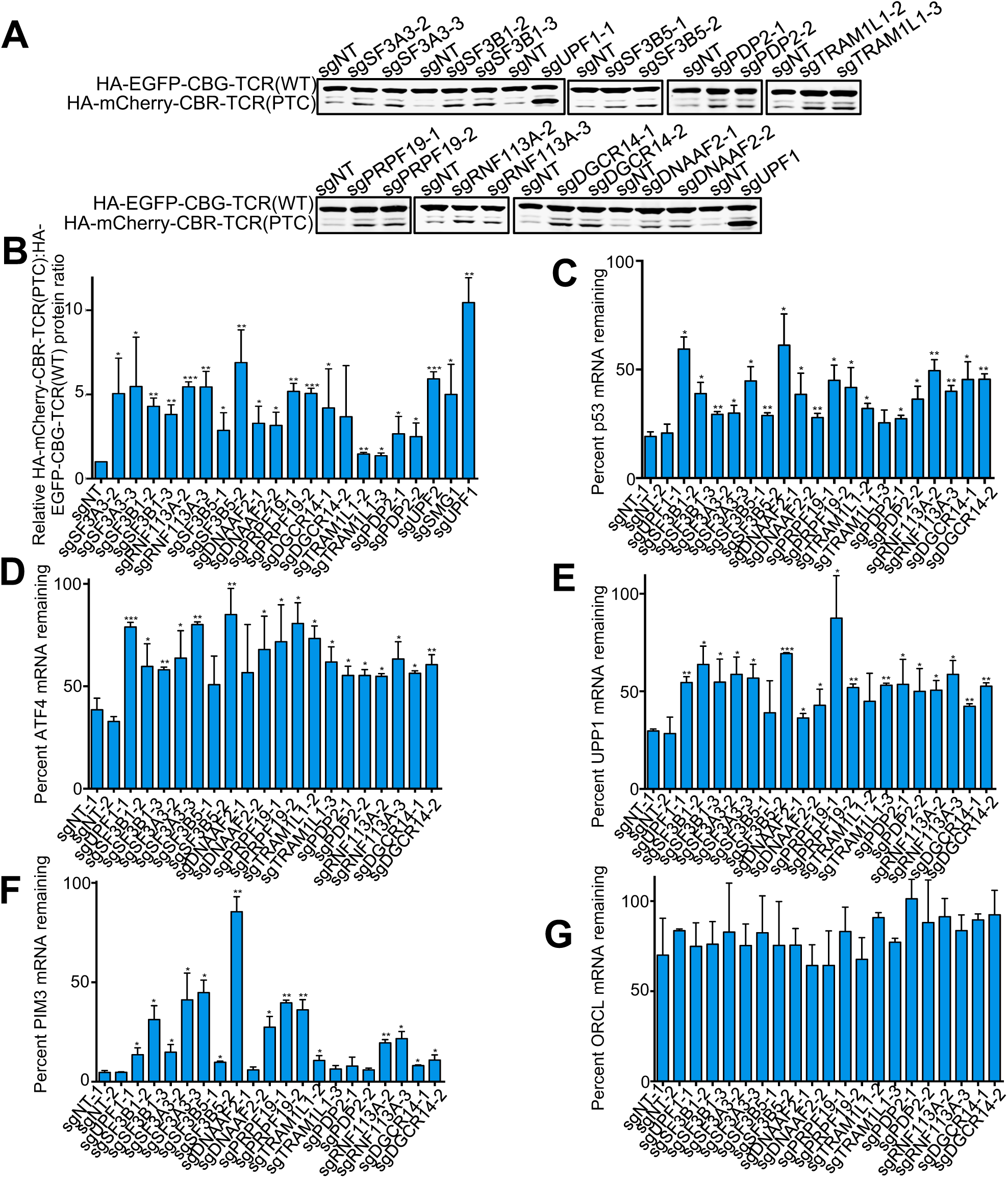
A. Western blot analysis of the protein products of the NMD reporter in Cas9-expressing U2OS reporter cells after sgRNA-mediated depletion of nine top-ranked genes individually. Two sgRNAs that are distinct from that in the original GeCKOv2 library were used for knockdown. B. Quantified results of the samples depicted in A. The ratios of the sgNT (nontargeting) control was normalized to 1. Data represent the mean ± SD of three independent experiments. ***p ≤ 0.001; **p ≤ 0.01; *p ≤ 0.05 (paired t-test). C. Effects of depletion of the 9 top-ranked genes individually on the stability of endogenous PTC-containing p53 mRNA in Calu-6 cells. Cells were transfected with sgRNA-Cas9 expression constructs and incubated for 6 days, and then treated with actinomycin D for 6 hours to inhibit transcription. Total mRNA was collected before and after actinomycin D treatment. p53 mRNA levels were measured using RT-qPCR. Data represent the mean ± SD of three independent experiments. *p ≤ 0.05; **p ≤ 0.01; (paired t-test). D-G. Effects of depletion of the 9 top-ranked genes individually on the stability of physiological NMD targets *ATF4* (D), *UPP1* (E), and *PIM3* (F) in Calu-6 cells. *ORCL* (G) is non-NMD target control. Samples were generated as depicted in C. Data represent the mean ± SD of three independent experiments. ****p ≤ 0.0001; ***p ≤ 0.001; **p ≤ 0.01; *p ≤ 0.05 (paired t-test).

**Supplementary Figure 3.**
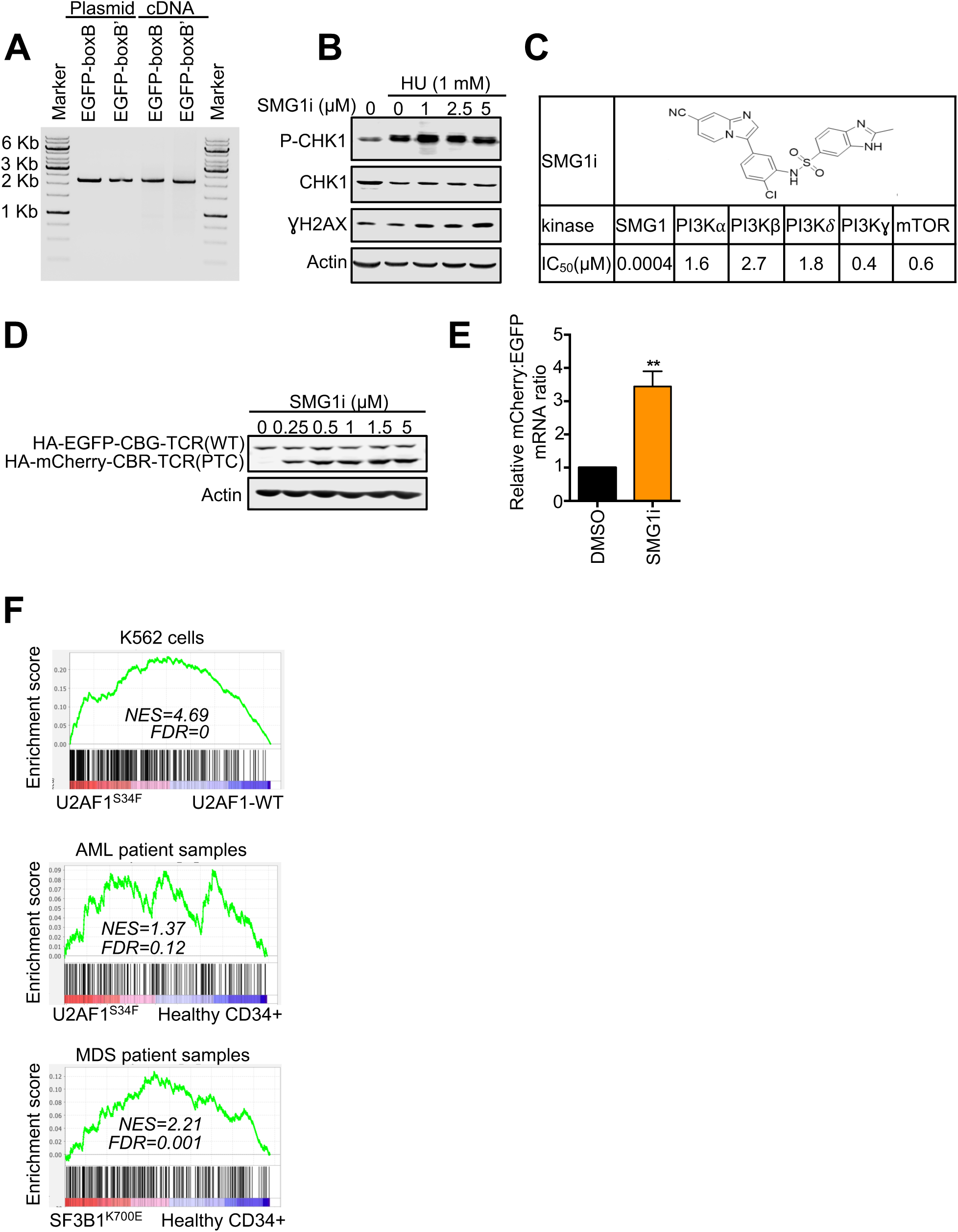
A. No cryptic splicing was detected in the intronless EGFP-boxB or EGFP-boxB’ tethering reporter RNA. Total RNA was extracted from U2OS cells expressing boxB or boxB’ reporter mRNAs, followed by RT using an oligo-dT primer. PCR was then used to generate cDNAs with primers that anneal to 5’UTR and 3’UTR of the reporter transcripts. B. SMG1i did not inhibit ATR activity towards CHK1 after replication stress. Western blot analysis of phosphor-CHK1 (pS345), total CHK1, and γH2AX in U2OS cells pre-treated with the indicated concentrations of SMG1i for 24 hours, and then treated with 1 μM hydroxyurea (HU) for 6 hours. C. SMG1i structure and its kinase inhibition activity against SMG1, and other PI3 kinase family members (PI3Kα, PI3Kβ, PI3Kγ, PI3Kδ, mTOR). D. Western blot analysis of the protein products of our new NMD reporter after treatment with SMG1i at indicated concentrations for 24 hours. E. Ratios of mCherry-containing reporter mRNA to EGFP-containing reporter mRNAs in U2OS reporter cells treated with 1 μM SMG1i for 24 hours. DMSO-treated cells were normalized to 1. Data represent the mean ± SD of three independent experiments. **p ≤ 0.01 (paired t-test). F. Gene Set Enrichment Analysis (GSEA) enrichment score plots for NMD target genes that are upregulated by SMG1i treatment. RNA was isolated from K562 cells (top), AML cells (middle), MDS cells (bottom), and control cells and sequenced to determine the effects of U2AF1^S34F^ expression (top and middle) and SF3B1^K700E^ expression (bottom, analysis of the data in Pellagatti et al.^121^) on gene expression. Individual genes in the gene set are represented by a black vertical bar at the bottom of the plot.

**Supplementary Figure 4.**
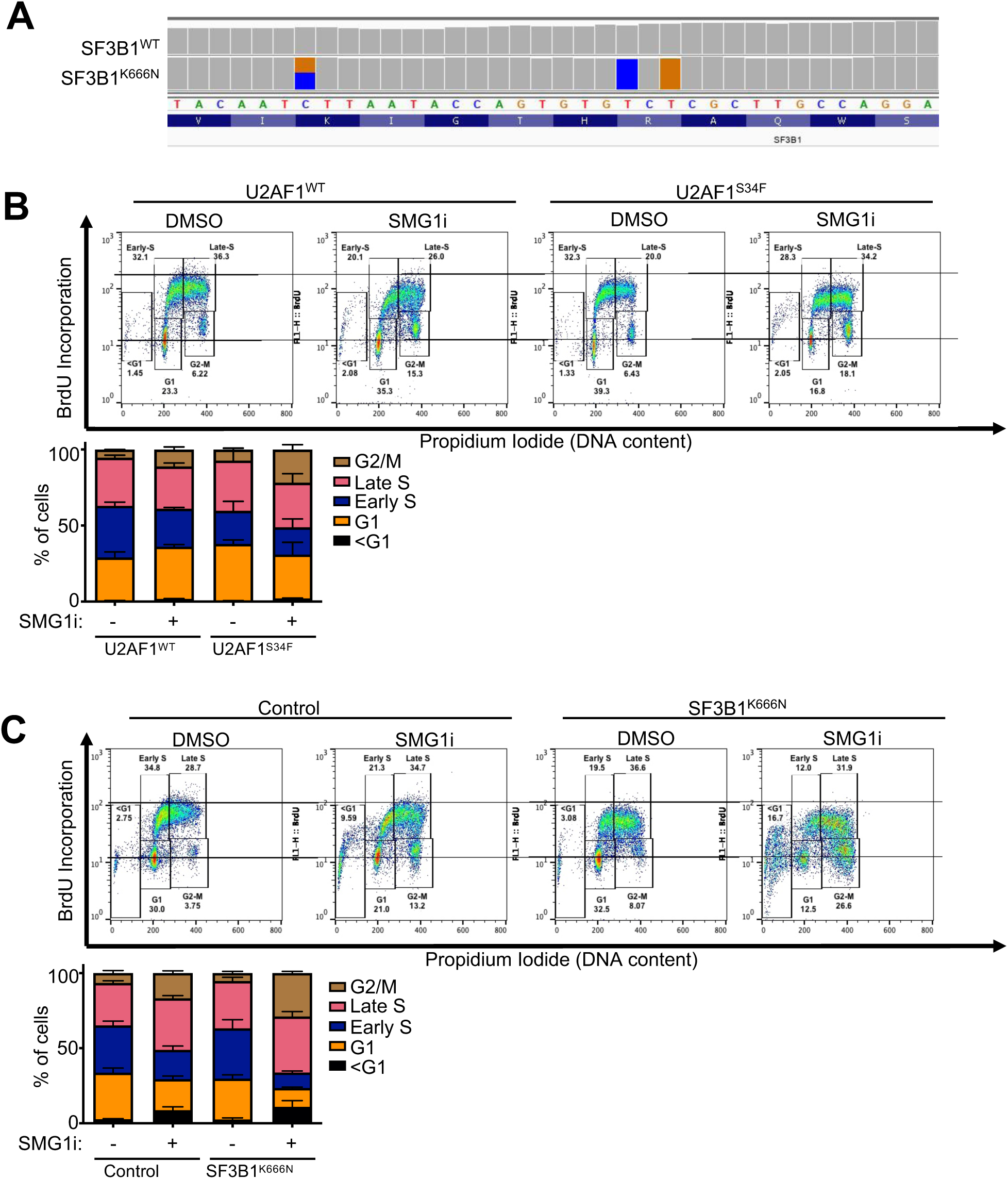
A. Genetically modified SF3B1^K666N^ K562 cells expresses 50% SF3B1^WT^ mRNA and 50% SF3B1^K666N^ mRNA. Sequence fragment density reads from RNA sequencing of K562 cells with or without SF3B1^K666N^ knock-in mutation aligned in IGV. The control cells express 100% SF3B1^WT^ mRNA. The knock-in cells express 50% mRNA containing the C to G mutation at position 1998 (which changes K at codon 666 to N, colored bars on the left) and 100% mRNA containing the blocking modifications (which do not change the amino acid) to prevent additional editing by Cas9 (colored bars on the right). B. Upper, effects of SMG1i treatment (1 μM, 3 days) on the cell cycle and DNA replication of K562 cells expressing U2AF1^WT^ or U2AF1^S34F^. Treated cells were pulsed labeled with BrdU for 30 min before being harvested for propidium iodide staining and flow cytometry. Lower, percentages of cells in different cell cycle phases after SMG1i treatment. Data represent the mean ± S.E.M. of three independent experiments. C. Upper, effects of SMG1i treatment (1 μM, 3 days) on the cell cycle and DNA replication of K562 cells with or without SF3B1^K666N^ knock-in mutation. Treated cells were pulsed labeled with BrdU for 30 min before being harvested for propidium iodide staining and flow cytometry. Lower, percentages of cells in different cell cycle phases after SMG1i treatment. Data represent the mean ± S.E.M. of three independent experiments.

**Supplementary Figure 5.**
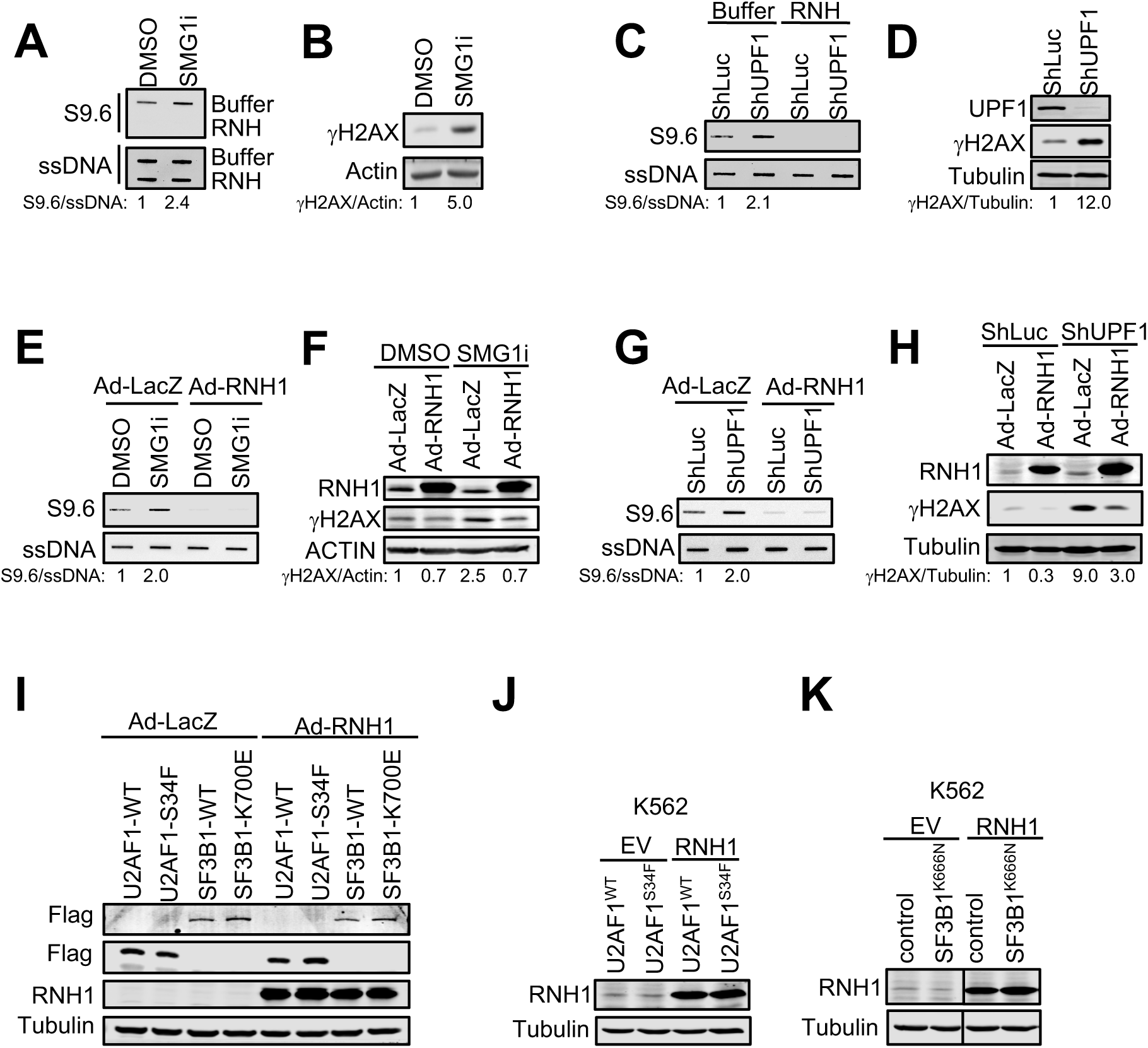
A. Effects of SMG1i treatment on R-loop levels. U2OS cells were treated with SMG1i (5 μM, 24 hours) followed by genomic DNA extraction for R-loop analysis. Total genomic DNA with or without RNase H (RNH) digestion were slotted on a membrane and S9.6 antibody was used to detect R-loops via slot blotting. ssDNA signal from denatured total genomic DNA was used as an input control. B. Effects of SMG1i treatment (5 μM, 24 hours) on γH2AX in U2OS cells. Samples were generated as described in A. C. Effects of shRNA-mediated knockdown of UPF1 on R-loop levels in U2OS cells. R-loops were detected via slot blotting in isolated total genomic DNA with or without RNase H (RNH) digestion D. Effects of shRNA-mediated knockdown of UPF1 on γH2AX in U2OS cells. Samples were generated as described in C. E. Effects of RNH1 expression on R-loops in SMG1i-treated U2OS cells. U2OS infected with adenovirus expressing lacZ control or RNH1 for 48 hours were treated with SMG1i (5 μM) for 24 hours. R-loops were detected by slot blotting. F. Effects of RNH1 expression on γH2AX in SMG1i-treated U2OS cells. Samples were generated as described in E. G. Effects of RNH1 expression on R-loop levels in UPF1-depleted U2OS cells. U2OS cells infected with lentivirus expressing shLuc or shUPF1 were infected with adenovirus expressing lacZ control or RNH1. H. Effects of RNH1 expression on γH2AX in UPF1-depleted U2OS cells. Samples were generated as described in G. I. Western blot analysis of RNH1, Flag-U2AF1 and Flag-SF3B1 in U2OS cells expressing U2AF1^WT^, U2AF1^S34F^, SF3B1^WT^, or SF3B1^K700E^ that were infected with adenovirus expressing lacZ or RNH1. J. Western blot analysis of RNH1 in K562 cells expressing inducible U2AF1^WT^ or U2AF1^S34F^ that were infected with adenovirus expressing empty vector (EV) or RNH1. K. Western blot analysis of RNH1 in wild type or SF3B1^K666N^ knock-in K562 cells that were infected with adenovirus expressing empty vector (EV) or RNH1.

